# Microbial association networks reveal hidden keystone taxa and cross-kingdom interactions after dryland wildfires

**DOI:** 10.64898/2026.07.28.741306

**Authors:** Arik Joukhajian, M. Fabiola Pulido Barriga, Peter M. Homyak, Sydney I Glassman

## Abstract

How wildfires reorganize soil microbial interactions is a key knowledge gap, particularly for drylands that cover nearly 40% of Earth’s surface and face increasing wildfire frequency with global change. We compared bacterial, fungal, and cross-kingdom association networks across four timepoints from 2 weeks to 3 years post-fire in two California dryland systems: a high-intensity chaparral shrubland fire and a low-intensity Eastern Joshua tree desert fire, using nearly identical sampling designs and molecular workflows. Wildfire increased bacterial-fungal associations more than bacterial or fungal interactions in both systems, with burned plots consistently shifting toward cooperative over competitive associations. Bacterial-fungal interactions also increased in burned relative to unburned desert plots, suggesting fire promoted microbial associations in desert soils. Although microbial richness declined by up to 61% one year after chaparral wildfire but remained unchanged in the desert, network clustering declined in both systems, indicating reduced community resilience independent of richness loss. Pyrophilous bacteria, including *Massilia* and *Noviherbaspirillum,* emerged as keystone taxa after chaparral wildfire, while generalist bacteria and the putatively pyrophilous Pyronemataceae fungus *Pseudotricharina* structured desert burned networks. Cross-kingdom network analysis revealed shifts in post-fire microbiomes invisible to traditional diversity metrics, highlighting bacterial-fungal interactions and keystone taxa as drivers of dryland post-fire succession.

## Introduction

Drylands cover 40% of the Earth’s land surface and are projected to expand due to climate change (Zeng and Yoon 2009; Feng and Fu 2013; Huang et al. 2016).

Drylands are increasingly burned by wildfires, driven by altered rainfall patterns, temporary blooms of plants, and rising temperatures (Stanton et al. 2023). Soil microbial communities, which mediate nutrient cycling, plant interactions, and ecosystem recovery (Jacoby et al. 2017; Crowther et al. 2019), and can influence ecosystem processes like carbon storage and gas emissions (Bardgett et al. 2008), are particularly sensitive to wildfire disturbance (Pressler et al. 2019; Certini et al. 2021). Yet microbiome responses to dryland wildfires are underexplored (Pressler et al. 2019), despite dryland susceptibility to nutrient loss (Schlesinger et al. 1990; Ren et al. 2024). Beyond this, not all arid ecosystems experience fire in the same way (Pausas and Keeley 2021). Fire intensity, or the heat output of a fire, can increase in denser drylands, while fire severity, which reflects the biological impact, can be extreme even without high intensity (Keeley 2009). Fire intensity and severity strongly influence how fires impact soil microbiomes, with high-intensity fires often leading to substantial reductions in microbial biomass and richness (Dooley and Treseder 2012; Nelson et al. 2022; Pulido-Chavez et al. 2023; Caiafa et al. 2023), while low-intensity fires may leave these metrics largely unchanged (Vega-Cofre et al. 2023; Glassman et al. 2023; Zhang et al. 2025).

While we have learned much from examining impacts of fires of differing intensities and severities on metrics like microbial biomass, richness, and composition (Dooley and Treseder 2012; Pressler et al. 2017), network analysis can give insights into how microbial interactions alter succession post-fire. Indeed, network analysis has been used to show how environmental stress can destabilize microbial networks (Hernandez et al. 2021), how bacterial networks are more susceptible to drought than fungal networks (de Vries et al. 2018) and how networks can become more connected and complex during succession after old field abandonment (Morriën et al. 2017).

Association networks allow inference of positive (potential facilitation or co-habitation) (Faust et al. 2015) and negative (potential competition or niche exclusion) interactions between taxa (Hirano and Takemoto 2019). Taxa are plotted as nodes in a network with the shifts in their relative abundance across many samples plotted as positive or negative edges linking each pair of associated taxa. Cohesion, a ratio of negative versus positive associations between taxa, has been used to assess community stability after microbial response to stress or disturbance (Hernandez et al. 2021; Birch et al. 2025). Modularity quantifies how tightly connected clusters of microbes are within a network with higher modularity often implying higher resilience, or ability to recover to original state (de Vries et al. 2018; Matchado et al. 2021). A few studies have investigated impacts of fires on networks, either showing that bacterial correlations were significantly reduced after controlled burns of biocrusts (Aanderud et al. 2019), or that bacterial network properties did not shift post-fire (Pérez-Valera et al. 2017; Whitman et al. 2022). Another study found that 9 years after a forest fire, cross-kingdom networks had lower cohesion (i.e. an overabundance of position associations) in heavily burned regions (Birch et al. 2025). What remains unknown is how bacterial-fungal interactions, as opposed to within kingdom interactions, differ during post-fire succession, and whether examining these cross-kingdom networks can uncover community level changes that are undetected by traditional metrics that focus on richness and biomass.

Network analysis can also identify keystone species (Jordán 2009), which are taxa with a greater impact on the community than their abundance reflects (Power et al. 1996). In microbial ecology, keystones are thought of as low-abundance species with a stabilizing effect on the community (Amit and Bashan 2023; Kajihara and Hynson 2024) that may drive microbiome structure (Banerjee et al. 2018), plant chemistry (Wang et al. 2023), or ecosystem multifunctionality (Luo et al. 2025). Network metrics such as high node degree (or number of connections per taxon) and betweenness centrality (taxon that lies on the shortest path between other taxa) (Kajihara and Hynson 2024) can generate hypotheses of keystone microbial taxa that can be tested for disproportionate impacts on microbiomes experimentally once isolates are cultured (Rawstern et al. 2025). Despite differing impacts on soil microbiomes, both high- and low-intensity fires can still promote the emergence of pyrophilous or fire-loving taxa, which are rare or absent pre-fire but bloom post-fire (Fox et al. 2022). Bacteria such as *Noviherbaspirillum* and *Massilia*, and fungi such as *Aspergillus*, *Coniochaeta*, *Pyronema*, and others have been isolated post-fire across the globe (Warcup 1990; Belmok et al. 2019; Enright et al. 2022, 2026; Soria et al. 2023; Li et al. 2024). Whether pyrophilous microbes act as keystone taxa, and whether those interactions are cross-kingdom, remains a knowledge gap.

Further, the impacts of wildfire on keystone taxa and their interactions, and network metrics like cohesion and modularity, can change over time. Microbial taxa may take up to 2 decades to recover from fire (Pérez-Valera et al. 2018), but reductions in microbial richness and biomass are often mitigated over time (Pulido-Chavez 2023; Fernández-González et al. 2023; Caiafa et al. 2023). Pyrophilous microbes also change in abundance over time (Pulido-Chavez et al. 2023), and some species like *Massilia* and *Blastococcus* can persist for up to 10 years post-fire (Fernández-González et al. 2023; Caiafa et al. 2023). Furthermore, trends over time can differ between bacteria and fungi in the same wildfire systems (Whitman et al. 2019, 2025; Pulido-Chavez et al. 2023; Caiafa et al. 2023; Yang et al. 2024), while other studies show that network complexity of both bacteria and fungi drop with reductions in connections between taxa immediately post-fire (Su et al. 2022). By comparing burned and unburned communities, it may be possible to highlight how post-fire microbial succession can shape taxa through associations even in cases without major shifts in biomass or dominance.

Here, we leverage datasets collected following the same experimental design, soil sampling and molecular methods, both sampling wildfires at numerous timepoints ranging from 2 weeks to 3 years post-fire in two drylands of differing fire intensity but both with high aboveground severity. The Holy Fire, which burned chaparral shrublands at high intensity, decreased richness of bacteria by 23% and fungi by 61% in the first year post-fire (Pulido-Chavez et al. 2023), whereas the Dome Fire, which burned Eastern Joshua tree desert at low intensity, did not affect bacterial or fungal richness (Joukhajian et al. 2026). Despite differing impacts on richness and composition, pyrophilous bacteria such as *Noviherbaspirillum*, *Tumebacillus*, *Paenibacillus*, *Massilia*, *Solirubrobacter*, and fungi such as *Aspergillus*, *Coniochaeta*, *Penicillium*, and *Naganishia* bloomed after both wildfires. We characterized microbial association networks in two California dryland ecosystems, an Eastern Joshua tree desert scrub and a chaparral shrubland, across 4 timepoints ranging from 2 weeks to 3 years post-fire, and asked: 1) how do bacterial, fungal, and bacterial-fungal associations and network metrics such as cohesion and modularity change? 2) can we identify keystone taxa, and how do they relate to pyrophilous microbes identified in these systems?

Overall, based on previous observations that the chaparral fire drastically reduced microbial richness (Pulido-Chavez et al. 2023), whereas the desert fire did not (Joukhajian et al. 2026), we expected to detect stronger impacts on network associations in the chaparral vs the desert fire. We predicted that cohesion would recover after the desert fire but not the chaparral fire after 3 years, but that both fires would result in long-term shifts in modularity, as compositional shifts alter associations. Since pyrophilous microbes were so dominant after the chaparral wildfire, we expected that they would emerge as keystone taxa following the chaparral but not the desert wildfire.

## Experimental Procedures

### Site descriptions

Our chaparral site was located in the Cleveland National Forest in California (Fig. 1), USA (33.678889, -117.516667) (Pulido-Chavez et al. 2023). The Holy Fire burned 94 km^2^ from August 6 to September 13, 2018 (Fig. 1C), primarily burning chaparral shrubland dominated by manzanita (*Arctostaphylos glandulosa*) and chamise (*Adenostoma fasciculatum*). Our desert site was in Mojave National Preserve in California, USA, centered around the Cima Dome (35.287598, -115.585376) (Joukhajian et al. 2026). The Dome Fire burned 175km^2^ around the Cima Dome (Fig. 1D) from August 15 to 24, 2020, killing 1 million Eastern Joshua trees (*Yucca jaegeriana*), and other perennial shrubs and grasses (Kaiser and Hughson 2020). While pockets of unburned trees remained present within the burn perimeter, tree death was 80% across our select burned plots. Despite the high Eastern Joshua tree mortality in the Dome Fire and high shrub mortality in the Holy Fire, leading to high aboveground severity for both, the fires burned at differing intensities, as reflected by average ash depth, which was 12.5X higher in Holy versus Dome Fire, which differed from mean ± standard error (SE) ash depth in cm of 4.63 ± 0.55 for Holy and 0.37 ± 0.07 for Dome Fire (Fig. 1E). Soils in the chaparral control plots are classified as Lithic Haploxerolls and mapped in the Friant series, as are soils in half of the plots within the burn scar, with the remaining burned plots on Typic Xerorthents, mapped in the Cieneba series.

**Figure 1.**
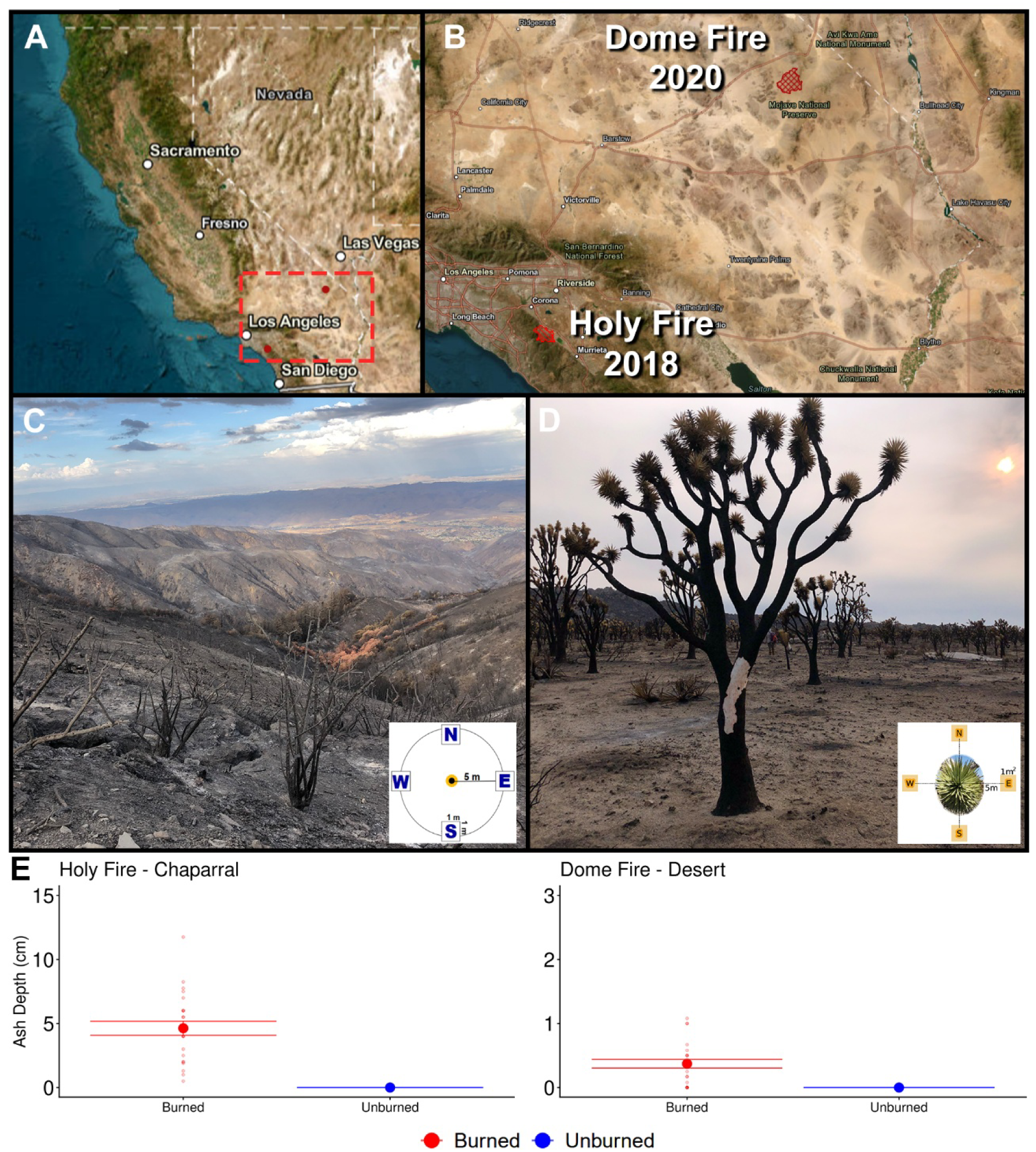
A) Overview of the sampling locations in California, with the red dash box capturing both wildfire sites. B) Fire perimeters (in red) of the 2018 Holy Fire and the 2020 Dome Fire in Southern California. C) The Holy Fire burn scar 17 days postfire and a subplot sampling diagram of one of our 9 plots. D) The Dome Fire burn scar showing a dead Eastern Joshua tree covered in *Neurospora discreta* 17 days postfire with a subplot sampling diagram of one of our 9 plots. E) Mean ash depth in centimeters at 17 days post-fire measured at the Holy Fire and the Dome Fire sites. Dots indicate the mean and bars indicate standard error (burned n=24, unburned n=12).

Both series are shallow sandy and gravelly loams (Soil Survey Staff 2024). The desert plots contain deep, well-drained, fine-loamy alkaline soils in the Minehart series (Soil Survey Staff 2015).

### Sample collection

Soon after each fire was extinguished (∼2 weeks), we established 9 plots (6 burned, 3 unburned) with four 1m^2^ subplots each in the cardinal directions 5m away from the plot centers (Fig. 1). From each of the 1m^2^ subplots, we collected 3-500g of soil using releasable bulb planters cleaned with 70% ethanol between subplots from the top 10cm of soil beneath organic or ash layer post-fire at 0.5, 8, 12, and 36 months in the desert scrub compared to 0.5, 6, 12, and 40 months in the chaparral shrubland.

We returned soil samples to the University of California Riverside (UCR) within 24-48 hours, immediately homogenizing and sieving (2mm) samples with ethanol-cleaned sieves, measured soil gravimetric moisture, and performed the KCl extraction method on fresh soil (Carter and Gregorich 2007) using previously reported methods (Joukhajian et al. 2026; Stephens et al. 2026). We stored KCl extracts at -20°C until we sent them to the UCR Environmental Science Research Lab to measure extractable ammonium, phosphate, and nitrate. We stored a subset of soil at -80°C for future molecular analysis and air-dried the rest. We measured pH with a VisionPlus pH6175 meter (Jenco Instruments Inc., San Diego, CA) after making a slurry of 10g of air-dried soil and 20mL of distilled water and shaking for one minute to mix.

### Molecular and bioinformatic work

Methods for DNA extraction, PCR amplification, and Illumina Miseq library prep for Holy Fire chaparral samples can be found in Pulido-Chavez et al. 2023 and for Dome Fire desert samples in Joukhajian et al. 2026. For chaparral samples we used Qiagen DNeasy PowerSoil kits, and for desert samples, we used Qiagen DNeasy PowerSoil Pro kits, due to discontinuation of the DNeasy PowerSoil kits. We amplified DNA in a two-step PCR using the ITS4-fun and 5.8S primer-pair to generate fungal ITS2 amplicons (Taylor et al. 2016), and the 515F and 806R primer-pair for archaeal and bacterial 16S amplicons (Caporaso et al. 2011).

Amplicons from 16S and ITS2 pools were combined at a 2:3 ratio to account for preferential sequencing of shorter amplicons, and sequenced together in Illumina Miseq 2x300 runs at the UCR Institute for Integrative Genome Biology. We used QIIME2 (Bolyen et al. 2019) to demultiplex and denoise reads, and used de-novo clustering at 97% to create Operational Taxonomic Units (OTUs) for genus-level comparisons, which are preferable for network analysis because environmental samples often lack species-level designations and intraspecific associations can inflate edge numbers (Röttjers and Faust 2018). We performed taxonomic assignment using the Qiime2 Naïve Bayes classifier with the SILVA (v138; Quast et al. 2013) and UNITE (v10; Kõljalg et al. 2005) databases. We uploaded sequences to the NCBI SRA under accessions PRJNA761539 for chaparral (Pulido-Chavez et al. 2023) and PRJNA1313334 for desert samples (Joukhajian et al. 2026).

### Network construction

We performed all analysis in R version 4.4.2 (R Core Team 2024). We constructed cross-kingdom and individual microbial association networks for bacteria and fungi using the SpiecEasi R package (Kurtz et al. 2015). For each timepoint, we subset 16S and ITS2 phyloseq objects to matching samples and by treatment prior to constructing networks (burned vs unburned). Since we only had 3 unburned plots (12 samples) compared to 6 burned plots (24 samples), to compare the effect of sample size on network properties, we created a second set of down-sampled burned networks (12 samples) to match the sample size of unburned networks (12 samples). For each kingdom, we collapsed OTUs to genus-level to avoid associations of OTUs matched to the same species, and then removed genera with fewer than 10 reads per treatment per timepoint (Jameson et al. 2023). Further, we only kept genera with a 16% prevalence per network by removing genera appearing in fewer than 2 samples in unburned networks or 4 samples in burned networks according to suggested best practices (Kajihara and Hynson 2024). After filtering, we used the “multi.spiec.easi” function with the Meinshausen–Bühlmann (MB) method (Meinshausen and Bühlmann 2006). We used 999 λ values (from a minimum λ ratio of 1e-3) selected via Stability Approach to Regularization Selection (StARS), with 300 repetitions and a selection threshold of 0.01. Feature tables used with the SpiecEasi package undergo centered-log ratio transformations to account for uneven sequence depth across samples and primer sets, allowing for cross-kingdom association network construction (Kurtz et al. 2015). This allows for conditional independence while drawing associations between nodes, such that each pair of nodes is evaluated independently from the rest of the network of genera.

### Network metrics

We used functions from the igraph package (Csardi 2013) to identify network metrics at each timepoint. To quantify community stability, we calculated abundance weighted mean cohesion per sample in each network. We extracted the signed edge weights from SPIEC-EASI networks by retrieving the adjacency matrix with the SpiecEasi function “getRefit” (Kurtz et al. 2015) and corresponding edge weight matrices with “symBeta”. We constructed undirected igraph objects from the adjacency matrix and assigned each edge its signed weight from the symmetrized beta matrix. We defined abundance-weighted cohesion as the ratio of summed negative edge weights to summed positive edge weights based on taxa present in each sample (Herren and McMahon 2017). We subset all edges connected to each present node (taxon) and summed edge weights for negative and positive separately, which we then used to generate a ratio per sample. We plotted mean cohesion per sample and tested significance of means using a linear model and showed pairwise significance between burned and unburned networks at each timepoint. We calculated modularity using the igraph package “modularity” after finding community structure with the function “cluster_louvain”. Higher modularity indicates that more edges connect nodes that are within the same module rather than across different modules (Csardi 2013).

### Keystones

We identified potential keystone taxa using two methods. First, we identified taxa with high betweenness centrality (Kajihara and Hynson 2024), indicative of nodes that frequently occur on the shortest path between other nodes. We labeled any node with >2 standard deviations of the mean betweenness per network, as well as those exhibiting >2 node degree, or connections to other nodes. We highlight these taxa in full network plots. Second, we calculated within-module connectivity (Zi) and among-module connectivity (Pi) adapted from (Deng et al. 2012) to classify genera into four categories: peripheral taxa with few connections (Zi < 2.5 and Pi < 0.62), module hubs with in-module connectivity or many edges (Zi > 2.5, Pi < 0.62), connector nodes with many edges across modules (Zi < 2.5, Pi > 0.62), and network hubs which are highly connected both within and across modules (Zi >2.5, Pi > 0.62). We used igraph to cluster modules from each network, which were subsets of treatment and timepoint, and used ggplot2 to combine results from all networks per site. We identified modules in which non-peripheral nodes appeared and plotted their positive and negative associations using the beta matrix based on weighted adjacency matrices per network. Although we use established methods in graph theory and microbial network analysis (Guimerà and Nunes Amaral 2005; Deng et al. 2012), we cannot confirm these taxa as true keystone species without further testing of their removal and the resulting consequences for the microbial community (Rawstern et al. 2025). However, we refer to these network-identified potential keystones hereafter as “keystones” for brevity.

## Results

### Fire impacts on ash, pH, extractable inorganic nitrogen, and microbial richness

Chaparral and desert wildfires burned at high versus low intensity as measured by variation in ash depth of 4.63 ± 0.6 cm in chaparral and 0.4 cm ± 0.07 in desert (Fig. 1E), which resulted in significantly increased pH in burned plots for both sites but with deserts being more basic than chaparral (Fig. S1A-B). Chaparral soil showed a significant increase in extractable ammonium and nitrate, although ammonium levels returned to unburned levels over time. In the desert, extractable ammonium levels remained elevated in burned plots (Fig. S1C-D), and nitrate increased in burned plots over time (Fig. S1E-F). Fire reduced bacterial and fungal richness at all time points from 0.5 to 40 months in chaparral but did not reduce microbial richness in desert (Fig. S2).

### Chaparral network statistics

In the chaparral, unburned networks were highly connected throughout succession, with the greatest number of nodes and edges across 16S, ITS2, and cross-kingdom networks (unburned nodes: 223-668; unburned edges: 36-576; burned nodes: 80-358; burned edges: 3-146; Table S1). The 16S and ITS2 networks from unburned plots peaked in associations within the first six months (16S: 79-122 edges; ITS2: 66-60 edges), while burned cross-kingdom networks had the most edges at 40 months (146). Despite some recovery by the final timepoint, burned cross-kingdom networks remained less connected than unburned ones (unburned maximum edges: 576; burned maximum edges: 146). Unburned chaparral networks also maintained consistently higher modularity than burned networks across all network types and timepoints, with burned modularity recovering only partially by 40 months (Fig. 2A). Cohesion, or the ratio of negative to positive edges, was significantly higher in unburned plots than in burned plots at all time points (Fig. 2C). Burned plots generally exhibited lower cohesion, resulting in networks with putatively lower levels of competition and redundancy and higher proportions of positive associations between taxa in single and cross-kingdom chaparral networks (Fig. S3A). This pattern was still apparent when subsampling burned networks to equivalent sample numbers of the unburned networks (from n=24 to n=12) in the chaparral (Fig. S4A).

**Figure 2.**
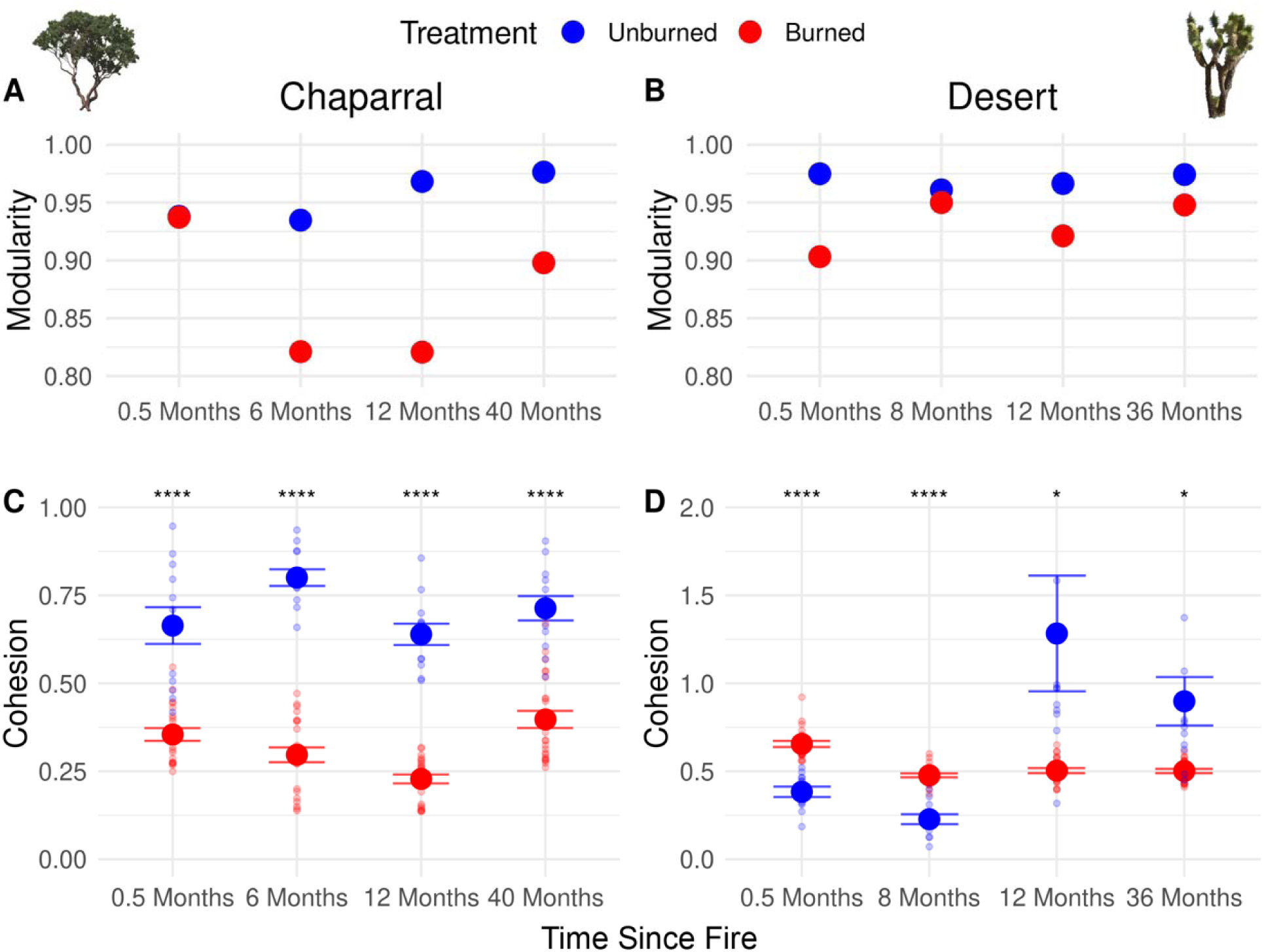
Network metrics from the chaparral (Holy Fire) and desert (Dome Fire) indicated by the dominant plants in those systems (Manzanita in chaparral and Eastern Joshua tree in Desert). Modularity is a measure of how often edges connect nodes within modules rather than across different modules, with one value per network (no standard error). Cohesion is a measure of the ratio of negative to positive edges, weighted by abundance of the nodes between the edges. Asterisks indicate significant pairwise t tests (ns = p ≥ 0.05, * = p < 0.05, ** = p < 0.01, *** = p < 0.001, and **** = p < 0.0001). Dots indicate the mean and bars indicate standard error (burned n=24, unburned n=12).

### Desert network statistics

Desert networks were overall smaller and less connected than those from the chaparral across all network types and timepoints. Unburned desert networks contained fewer nodes and edges than chaparral unburned networks (16S: 221-271 nodes, 18-40 edges; ITS2: 122-158 nodes, 8-17 edges; cross-kingdom: 349-429 nodes, 62-117 edges), and edge counts generally declined from 0.5 to 36 months (Table S2). Burned desert networks had comparable or greater edge counts than unburned desert networks across all timepoints (16S: 57-74 vs. 18-40; ITS2: 15-28 vs. 8-17; cross-kingdom: 139-170 vs. 62-117), suggesting fire promoted more microbial associations in desert soils than would otherwise occur. Despite this, unburned desert networks maintained higher modularity than burned networks across all timepoints and network types (Fig. 2B). Unlike the chaparral, cohesion in the desert was significantly higher in the burned than unburned plots at the first two timepoints (Fig. 2D), but the trend reversed and matched the chaparral with higher cohesion in unburned than burned plots after the first year. Subsampling desert burned networks to match unburned networks (n=24 to n=12) eliminated statistical differences in all timepoints except at 8 months, where burned networks still had higher cohesion than unburned networks (Fig. S4B).

### Single and cross-kingdom networks

Node and edge numbers differed among bacterial, fungal, and cross-kingdom networks. In both sites, bacterial networks contained more nodes (taxa) and edges (associations) than fungal networks at each timepoint. In the chaparral, unburned bacterial networks ranged from 289-391 nodes and 49-122 edges, while unburned fungal networks ranged from 223-285 nodes and 36-66 edges (Table S1). In the desert, unburned bacterial networks ranged from 221-271 nodes and 18-40 edges, compared to unburned fungal networks of 122-158 nodes and 8-17 edges (Table S2). Cross-kingdom networks consistently contained the largest node sets and substantially more edges than either single-kingdom network type, reflecting the addition of associations between archaeal, bacterial, and fungal taxa. This pattern was most pronounced in unburned chaparral plots (Fig. 3), where combining datasets produced several hundred additional edges relative to either 16S or ITS2 networks alone (peaking at 668 nodes and 576 edges). In the desert, cross-kingdom networks also displayed higher edge counts than single-kingdom networks but with smaller overall node numbers (maximum cross-kingdom nodes and edges: 429 and 117; maximum single kingdom: 265 and 74; Fig. 4).

**Figure 3.**
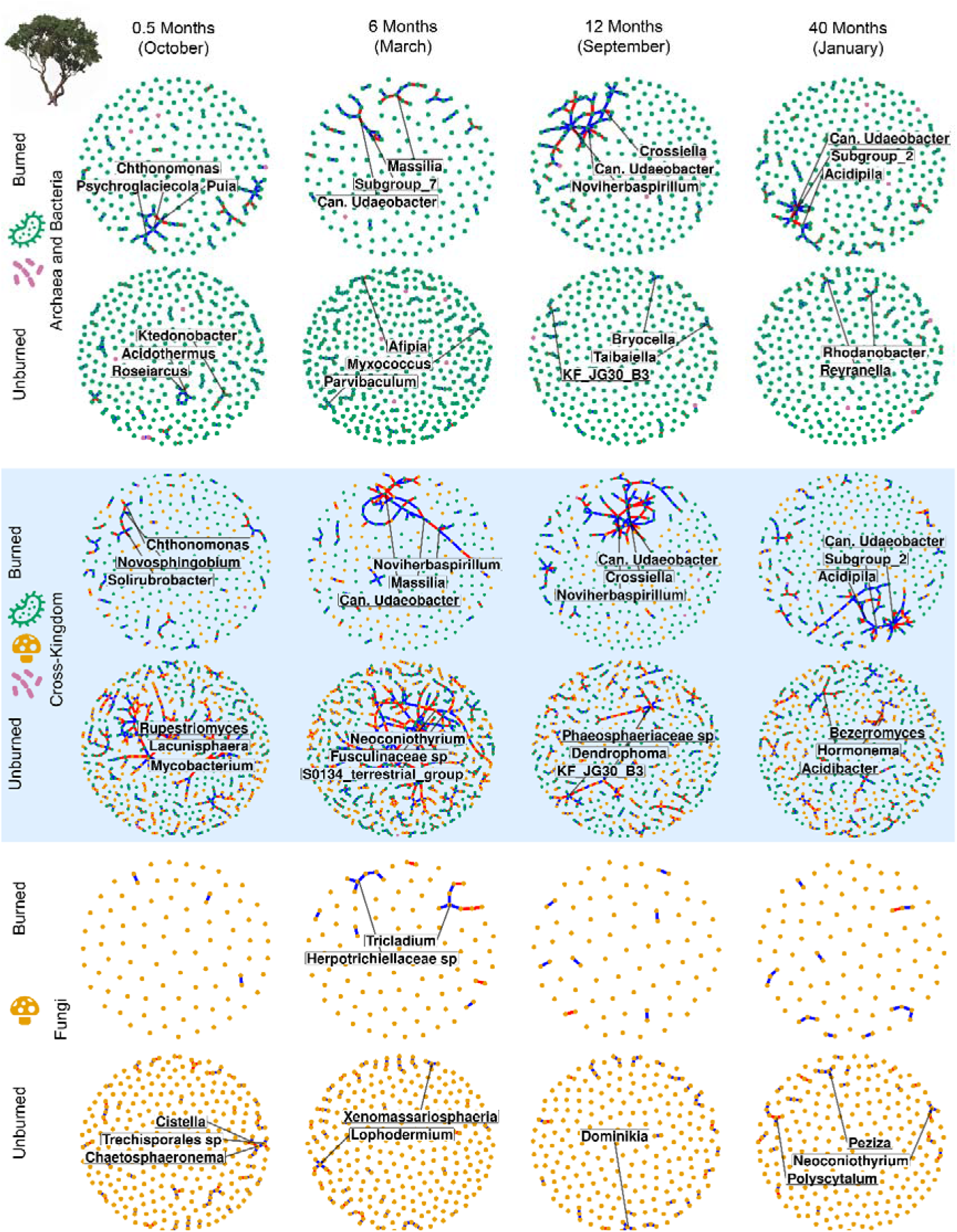
Networks from chaparral 16S and ITS2 sequence data in purple. Cross-kingdom networks displayed in blue. Nodes represent genera and edges represent inferred conditional associations, with positive associations in blue and negative in red. We retained genera if they had ≥10 reads and ≥16% prevalence within each network. All edges shown are those retained by StARS model selection. Node colors correspond to kingdoms, with pink being archaea, green being bacteria, and orange being fungi. The top 3 taxa with >2 std. dev. betweenness and >2 edges are labelled per network.

**Figure 4.**
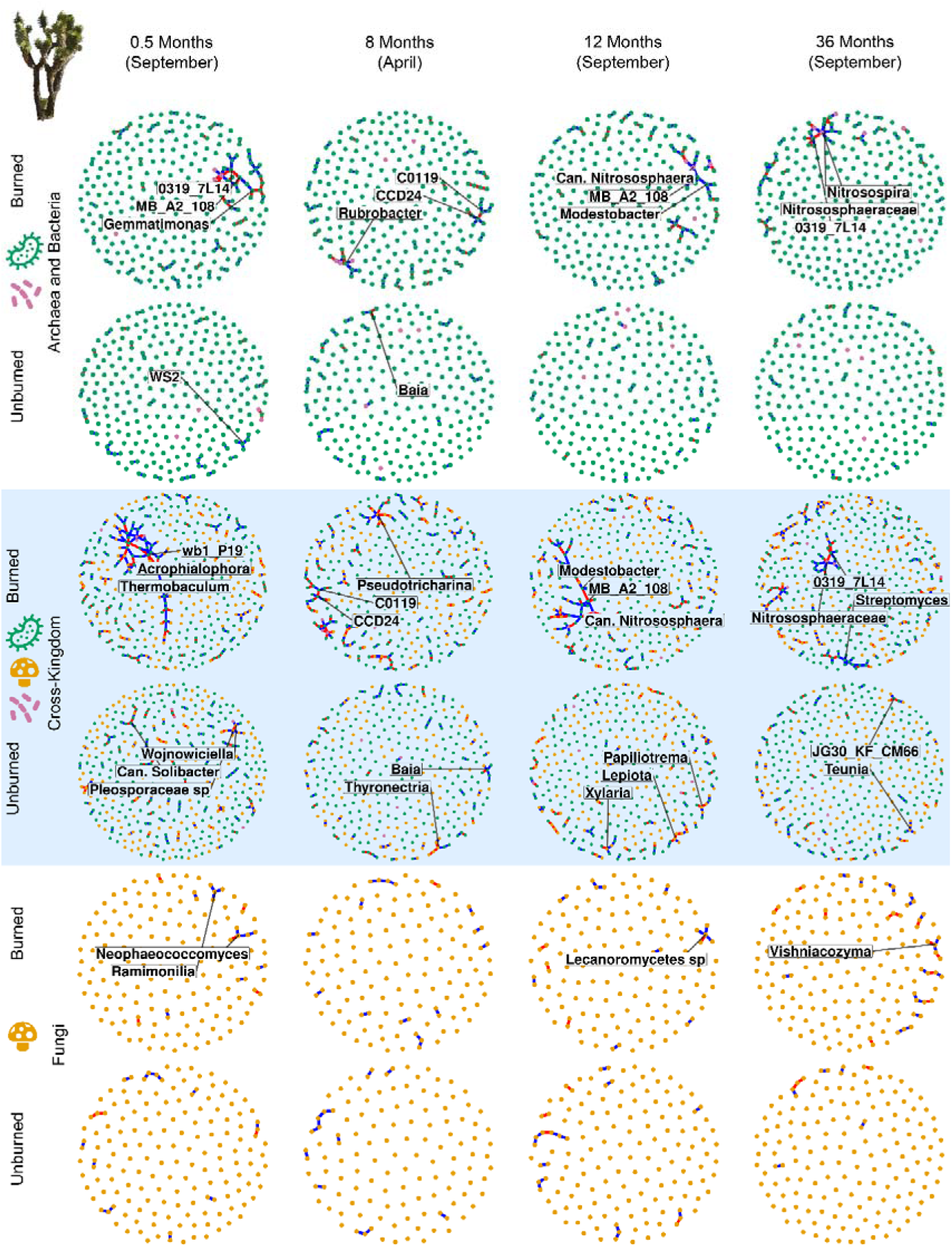
Networks from desert 16S and ITS2 sequence data displayed on white. Cross-kingdom networks displayed in blue. Nodes represent genera and edges represent inferred conditional associations, with positive associations in blue and negative in red. We retained genera if they had ≥10 reads and ≥16% prevalence within each network. All edges shown are those retained by StARS model selection. Node colors correspond to kingdoms, with pink being archaea, green being bacteria, and orange being fungi. The top 3 taxa with >2 std. dev. betweenness and >2 edges are labelled per network.

### Keystones

In each network, we identified putative keystone taxa based on within-network properties (>2 standard deviation betweenness, and >2 edges). Single-kingdom keystones were rare using these thresholds, especially for fungal networks. In desert cross-kingdom networks, these keystones were also sparse, particularly in unburned networks. Unburned chaparral network keystones were abundant, and still present after applying Zi-Pi thresholds for better cross-network comparisons (Table S3). Burned cross-kingdom network chaparral keystones often co-associated, appearing in the same modules (Fig. 3, 6). We classified keystones using within-module degree (Zi) and among-module participation coefficient (Pi) scores based on the presence of edges and presence of taxa within modules, with cutoffs using established thresholds (Guimerà and Nunes Amaral 2005), visualized in Figs. 5A for chaparral and 5B for desert. Taxa in the tan region (Zi > 2.5, Pi < 0.62) are module hubs, highly connected within their own module but not broadly connected across modules, whereas taxa in the green region (Pi > 0.62) are connectors or network hubs, linking taxa across different modules regardless of within-module connectivity. Peripheral taxa in the white region (Zi < 2.5, Pi < 0.62) have few connections within or across modules and are not considered keystones. This distinction matters ecologically, as module hubs may stabilize local clusters of co-occurring taxa, while connectors may mediate broader community-level associations across functionally distinct groups (Banerjee et al. 2018; Kajihara and Hynson 2024). Although taxa from unburned networks did meet these thresholds as connector and module hub taxa in the chaparral (Fig. S5, Table S3), we left them unlabeled (Fig. 5A) and focus on burned network keystones here.

**Figure 5.**
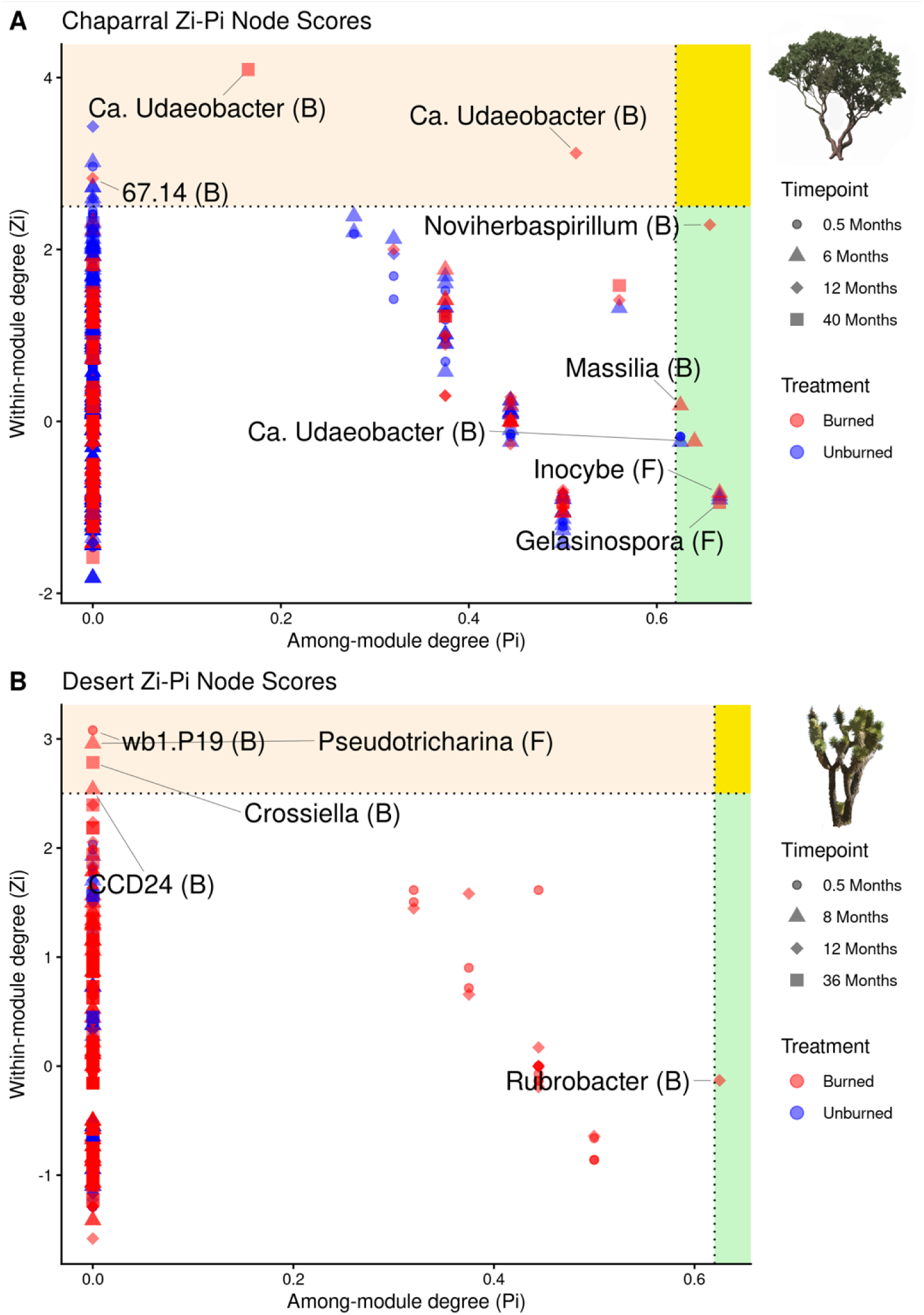
Zi-Pi plots of cross-kingdom networks from the A) chaparral and B) desert sites. Only burned network taxa are labeled, and kingdom is indicated by (B) for bacteria and (F) for fungi. Shapes indicate timepoints, and colors indicate treatment (red for burned and blue for unburned). Peripheral taxa in the white region are below the threshold of being module hubs or connector nodes. Peripheral taxa have few connections (Zi < 2.5 and Pi < 0.62), module hubs have in-module connectivity or many edges (Zi > 2.5, Pi < 0.62; **tan region**), connector nodes have many edges across modules (Zi < 2.5, Pi > 0.62; **green region**), and network hubs are highly connected both within and across modules (Zi >2.5, Pi > 0.62; intersection; **yellow region**).

### Chaparral keystone taxa

We examined whether genera from burned chaparral plots emerged as keystones as either connectors with high among-module connectivity, or module hubs with high within-module connectivity (labeled taxa, Fig. 5A). *Massilia* (phylum Pseudomonadota, connector; Module 1), *Candidatus Udaeobacter* (phylum Verrucomicrobia, hub; Module 2), and the ectomycorrhizal fungus *Inocybe* (Basidiomycota, hub; Module 1) were keystones at 6 months, *Noviherbaspirillum* (phylum Pseudomonadota; Module 10), *Candidatus Udaeobacter* (Module 11), and a bacterium from *67-14* (phylum Actinomycetota; Module 33) were module hubs at 12 months, and *Candidatus Udaeobacter* (hub, Module 16) and the fungus *Gelasinospora* (Ascomycota, connector; Module 21) were keystones at 40 months post-fire (Fig. 6).

**Figure 6.**
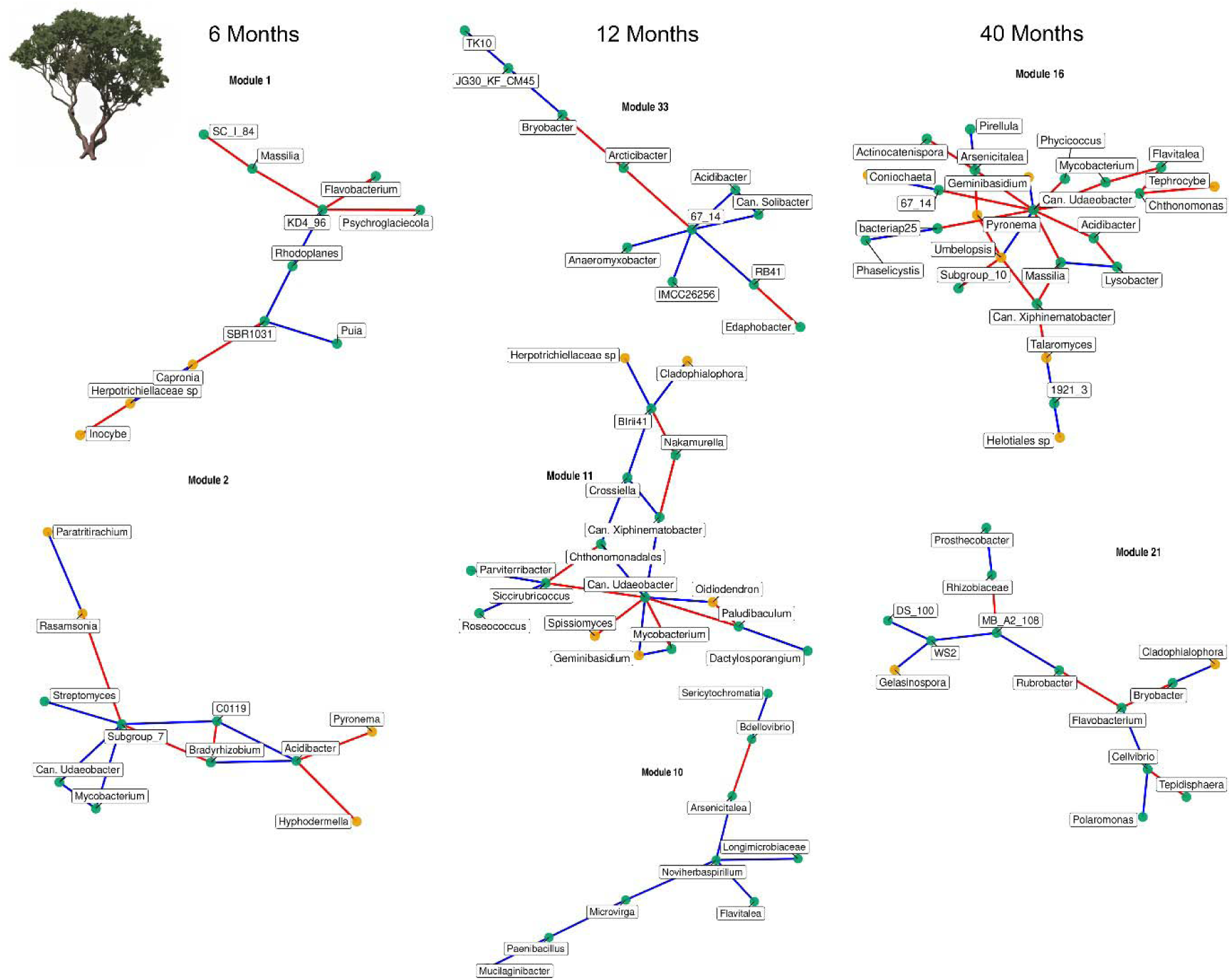
Modules of burned network hubs in the chaparral site. Modules 1 and 2 were at 6 months, 11 and 33 were at 12 months, and 16 and 21 were at 40 months post-fire. Blue edges are positive associations and red edges are negative. Node colors correspond to kingdoms, with pink being archaea, green being bacteria, and orange being fungi.

While some modules containing these keystone taxa were exclusively bacterial, others had many bacterial-fungal associations. For example, at 40 months, the Verrucomicrobia *Candidatus Udaeobacter* (Module 16) associated with 3 Pseudomonadota: *Arsenicitalea, Massilia, Acidibacter,* and the fungi *Coniochaeta* (Ascomycota) and *Geminibasidium* (Basidiomycota), while *Gelasinospora* (Ascomycota, connector; Module 21) shared a module with bacteria from *Rubrobacter* (Actinomycetota) and *Bryobacter* (Acidobacteriota) and another Ascomycota fungus *Cladophialophora*. Notably, *Candidatus Udaeobacter* was the only taxon appearing as a hub across three timepoints, and *Massilia* shifted from a hub role at 6 months to a peripheral associate of *Candidatus Udaeobacter* by 40 months, suggesting a successional transition in the importance of these pyrophilous taxa over time. Unburned networks had more keystone taxa than burned networks in the chaparral (13 vs 8; Fig. S5). Notably, *Mycobacterium* (Actinomycetota) is a connector hub (Table S3) in unburned networks, yet is found in the burned modules positively associated with *Candidatus Udaeobacter* at 6 months and negatively associated with it 40 months. No other unburned keystone taxa from the chaparral reappeared in burned network modules.

*Desert keystone taxa:* In contrast to chaparral, we only detected keystone taxa from burned networks in the desert (Fig. 5B). These included the bacterium *wb1-P19* (Pseudomonadota, module hub) at 0.5 months, which was centrally connected to *bacteriap25* (Pseudomonadota), and *Modestobacter* (Actinomycetota), alongside several fungal associates including *Glomeraceae* sp. (Mucoromycota) and the Ascomyota *Acrophialophora,* Pyronemataceae sp., and *Spissiomyces*, indicating cross-kingdom associations from the earliest post-fire timepoint (Module 1; Fig. 7). At 8 months, the fungus *Pseudotricharina* within the family Pyronemataceae (Ascomycota, hub; Module 8) was connected to the Ascomycota genera *Neophaeococcomyces* and *Inopinatum* alongside the bacterium *Pseudonocardia* (Actinomycetota), while CCD24 (Chloroflexota, hub; Module 9) associated with the bacterium *Segetibacter* (Pseudomonadota), *Polycyclovorans* (Pseudomonadota), and *Roseimicrobium* (Verrucomicrobia). At 12 months, *Rubrobacter* (Actinomycetota, connector; Module 31) was in a module with *Gaiella* (Actinomycetota), *Blastococcus* (Actinomycetota), the archaean *Can. Nitrososphaera* (Thermoproteota), and the fungus *Aspergillus* (Ascomycota). At 36 months, *Crossiella* (Actinomycetota, module hub; Module 57) was associated with an archaeon in Marine Group II (Thermoplasmatota), the fungi *Conocybe* (Basidiomycota), *Venturia* (Ascomycota), and *Phaeococcomyces* (Ascomycota), and the bacterium *Acidibacter* (Pseudomonadota), thus forming cross-kingdom associations. Across all timepoints, desert burned keystone modules contained fungal associates, although only the keystone *wb1-P19* had direct positive associations with bacteria and fungi immediately post-fire at 0.5 months, along with the keystone *Rubrobacter*, which positively associated with archaea and bacteria at 12 months.

**Figure 7.**
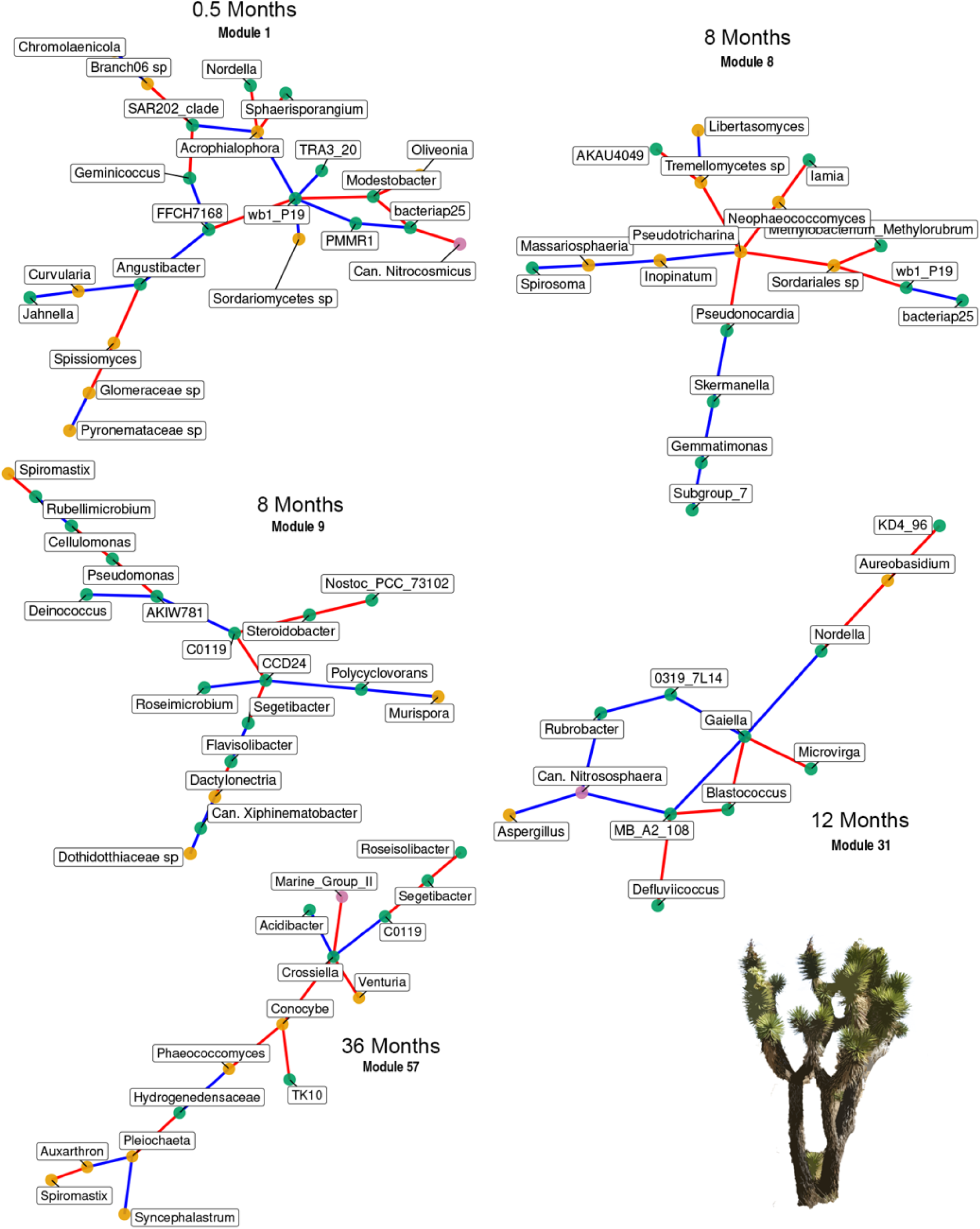
Modules of burned network hubs from the desert site. Module 1 was at 0.5 months, Module 8 and 9 were at 8 months, Module 31 was at 12 months, and Module 57 was at 36 months post-fire. Blue edges are positive associations and red edges are negative. Node colors correspond to kingdoms, with pink being archaea, green being bacteria, and orange being fungi.

## Discussion

We set out to determine how wildfire shapes bacterial, fungal, and bacterial-fungal associations and network metrics such as cohesion and modularity, and whether keystone taxa emerge and align with pyrophilous microbes identified in these systems. Overall, based on previous observations that the chaparral fire drastically reduced microbial richness (Pulido-Chavez et al. 2023), whereas the desert fire did not (Joukhajian et al. 2026), we expected to detect stronger impacts on network associations in the chaparral versus the desert fire. We predicted that cohesion would recover after the desert fire but not the chaparral fire after 3 years, but that both fires would result in long-term shifts in modularity, as compositional shifts alter associations. Since pyrophilous microbes were so dominant after the chaparral wildfire, we expected that they would emerge as keystone taxa following the chaparral but not the desert wildfire. In both wildfires, burned plots showed reduced modularity compared to unburned plots, though declines were strongest in chaparral networks. Wildfires appeared to reduce competition and increase positive interactions in both ecosystems, although the pattern was stronger and more persistent in chaparral, where fire intensity was higher. Keystone taxa, which were diverse in unburned plots in chaparral but nonexistent in unburned plots in desert, revealed interesting impacts of pyrophilous microbes in burned plots. For example, pyrophilous bacteria that have been detected across many ecosystems (Woolet et al. 2022; Whitman et al. 2022; Pulido-Chavez et al. 2023; Caiafa et al. 2023), such as *Massilia* and *Noviherbaspirillum*, were identified as keystones in burned plots in chaparral. In contrast, in the desert burned plots, keystone taxa included two bacteria in the phylum Actinomycetota, the oligotrophic *Crossiella* (Martin-Pozas et al. 2024) and stress-tolerant *Rubrobacter* (Ferreira et al. 1999), as well as the fungus *Pseudotricharina*, which is newly described but is a member of the pyrophilous family Pyronemataceae (Fox et al. 2022). Both *Rubrobacter* and *Crossiella* reappear in chaparral modules but are only keystones in the desert. Similarly, bacteria like *Acidibacter* and *Microvirga* and fungi like *Spissiomyces* and *Cladophialophora* reappear across modules in both systems without reaching keystone status in either.

Despite some taxa such as *Massilia* responding positively to fire in both systems, no keystone taxa overlapped between burned chaparral and desert networks, highlighting the habitat-specificity of keystone roles in post-fire succession.

### Network cohesion across time

Post-fire network cohesion revealed a long-term rise in positive microbial associations across both systems. This shift likely reflects the post-fire processes in which fire eliminates competitors and reduces community diversity, leaving surviving taxa in a less competitive environment where interactions are more probable than niche exclusion. Further, the remaining taxa may increasingly depend on one another for shared resource acquisition in the altered post-fire substrate environment, making communities more interdependent and potentially more vulnerable to future disturbance. Similar shifts toward more positive correlations under stress have been observed following drought and nutrient enrichment, where facilitative interactions temporarily enhance stability but may increase susceptibility to further disturbance (Coyte et al. 2015; de Vries et al. 2018). Similarly, a study of cross-kingdom networks nine years after a forest fire similarly found an overabundance of positive associations in heavily burned regions (Birch et al. 2025). Our findings extend this pattern to dryland wildfires across multiple timepoints, a context that has not previously been examined.

The rise in positive associations in burned plots may partly reflect cross-feeding among pyrophilous taxa, as genomic work has shown that pyrophilous bacteria carry genes for degrading pyrogenic organic matter (PyOM), and horizontal gene transfer of these pathways between pyrophilous bacteria and fungi has been documented (Sari et al. 2025, 2026). In the desert, the rise in positive associations occurred later, following rainfall that reactivated unburned soils, consistent with delayed microbial responses to moisture pulses in arid ecosystems (Blazewicz et al. 2014). Despite stable richness through time after the desert fire (Joukhajian et al. 2026), changes in cohesion revealed hidden structural sensitivity not captured by diversity metrics alone. In contrast, chaparral networks were initially fragmented after high-severity fire but slowly grew in node and edge density by 40 months, suggesting a slow reestablishment of microbial interactions similar to post-fire network recovery patterns in forest soils (Su et al. 2022; Woolet et al. 2022).

### Network modules across sites

Unburned chaparral networks were larger and more connected than burned networks across all network types, whereas desert networks were smaller overall with more comparable connectivity between burned and unburned plots. Unburned chaparral plots consistently exhibited higher node and edge numbers than burned plots across all timepoints, with dense module structure and high modularity. Unlike a study that found that experimental fire in Mediterranean shrubland altered community composition and assembly mechanisms without changing general network parameters (Pérez-Valera et al. 2017), chaparral wildfire in our study led to reductions in network size and connectivity, suggesting that wildfire severity or ecosystem context may determine whether post-fire filtering translates into detectable network-level changes. Yet by 40 months, burned chaparral networks approached unburned levels in both nodes and edges, which can reflect increased interactions (Jiang et al. 2022). These increases in associations imply a community adapted to high-return fire intervals, capable of rebuilding associations following prolonged organic matter availability. The decrease in modularity and cohesion in burned chaparral networks, as a result of increased positive edges, suggests shared niche preferences and sustained interdependence, likely among decomposers breaking down post-fire substrates (Coyte et al. 2015). In fact, many genes for PyOM degradation and N cycling were found to increase over time following the Holy Fire (Pulido Barriga et al. 2025).

While chaparral wildfire reduced modularity and cohesion, the lower intensity desert wildfire promoted increased connectivity with burned networks having more nodes and edges than unburned networks across our 3-year study period. This increase in connectivity may reflect that fire temporarily increases pH (Certini 2005), which is a deterministic abiotic filter that normally constrains bacterial network complexity in deserts (Dong et al. 2025). Thus, desert fires created conditions where stochastically assembled generalist taxa can form broader associations than would otherwise occur in undisturbed soil (Xu et al. 2022), such as the cross-kingdom associations we found post-fire.

### Network insights from cross-kingdom combinations

Bacterial-fungal cross-kingdom interactions emerged as more prominent than within kingdom interactions across both ecosystems, although interestingly, bacterial-fungal interactions were greater in burned plots than unburned throughout the desert successional timepoints but not in the chaparral. Furthermore, burned desert keystone modules contained fungal associates at every timepoint, from the bacterium *wb1-P19* associating with the fungi *Acrophialophora* and Sordariomycetes sp at 0.5 months to the bacterium *Crossiella* associating with the fungi *Conocybe* and *Venturia* at 36 months. This suggests that bacterial-fungal association was a consistent and persistent feature of post-fire desert network structure rather than an early successional artifact, potentially reflecting stress-induced metabolic cross-feeding between kingdoms (Velez et al. 2018; Amarnath et al. 2023) under the sustained abiotic pressures of post-fire desert soil.

The chaparral shubland showed a more muted cross-kingdom increase in burned relative to unburned plots, likely because severe richness losses reduced the pool of taxa available to form new cross-kingdom associations. Nevertheless, cross-kingdom associations were still present within burned chaparral modules, including the bacterium *Candidatus Udaeobacter* associating with the fungi *Pyronema* and *Geminibasidium* at 40 months, and the Ascomycota fungi *Gelasinospora* and *Cladophialophora* sharing a module with the Actinomycete bacteria *Rubrobacter* in Module 21 (Fig. 6). Although cross-kingdom associations are correlational and cannot confirm direct interactions (Guseva et al. 2022), the demonstrated introgression of a plasmid from *Noviherbaspirillum soli* carrying PyOM-breakdown genes into the pyrophilous fungus *Coniochaeta* (Sari et al. 2025, 2026) suggests that our association networks are detecting real interactions with important consequences for post-fire ecosystem regeneration. Furthermore, network analyses can generate testable hypotheses about bacterial-fungal functional relationships (Rawstern et al. 2025).

While very few studies have examined cross-kingdom networks in a wildfire setting, limiting our basis for comparison, one study in a high severity burned forest regions nine years post-fire found reduced cohesion in burned plots (Birch et al. 2025), consistent with our results in the high intensity chaparral fire. Studies that have analyzed within kingdom networks across fire chronosequences found that bacterial and fungal networks responded in opposite directions post-fire in subtropical forests such that bacterial became less complex and fungal networks increased in connections and modularity (Yang et al. 2024). In boreal forests, network complexity in both bacteria and fungi took 17 years post-fire to recover to that of unburned levels (Su et al. 2022). In non-fire systems, the cross-kingdom edge increase was detectable in a forested wetland study, which found a marginal increase from single-kingdom networks (332 and 319 edges in bacterial and fungal networks) to the cross-kingdom network (340 edges; Wu et al. 2023). Another study examining rice seed microbiomes identified substantial increases to edge and node counts after switching from single to cross-kingdom networks (Lee et al. 2022), where fungi acted as modular connectors of bacteria in this plant-microbe interface.

### Chaparral keystones

A central finding of this study is that pyrophilous microbes serve as keystone taxa following high-intensity chaparral wildfire, consistently dominating module hubs in burned but not unburned networks. This may reflect shared habitat preferences among pyrophilous taxa or associations with distinct functional modules of post-fire substrate degradation. At 6 months, the Pseudomonadota bacterial genus *Massilia*, which has been identified as pyrophilous across many ecosystems (Whitman et al. 2019, 2022; Enright et al. 2022; Soria et al. 2023; Fernández-González et al. 2023) appeared as a connector across different modules, reflecting its abundance in the community at that timepoint (Pulido-Chavez et al. 2023) and its broad metabolic capacity for post-fire substrate utilization. Genomic work has shown that *Massilia litorea* has rearranged naphthalene degradation and beta-ketoadipate pathways acquired via horizontal gene transfer from *Noviherbaspirillum soli* plasmids (Sari et al. 2025), suggesting that shared PyOM-degradation gene content between these two pyrophilous bacteria may support their co-occurrence in early post-fire networks. Wildfire has also been shown to shift the functional potential of *Massilia*-dominated communities toward aromatic C and nitrogen cycling in chaparral (Pulido Barriga et al. 2025) and forests (Nelson et al. 2024), consistent with its connector role linking taxa involved in post-fire substrate degradation across modules at 6 months.

Beyond *Massilia*, whose dominance in post-fire chaparral soils made it an unsurprising network hub, burned chaparral networks were also structured by *Inocybe* (Basidiomycota, module hub) and *Candidatus Udaeobacter* (Verrucomicrobia, module hub), taxa that would have been overlooked by conventional abundance-based approaches but emerged as keystones through network analysis. As an ectomycorrhizal fungus (EMF) emerging following widespread loss of other EMF taxa post-fire (Nelson et al. 2022; Caiafa et al. 2023), *Inocybe* likely bridges mycorrhizal plant seedlings and the broader microbial community across modules, consistent with the role of EMF in enhancing connectivity across multi-kingdom networks in forests (Yang et al. 2022).

*Candidatus Udaeobacter*, a Verrucomicrobia genus with a streamlined genome focused on key enzymatic pathways in amino acid synthesis (Brewer et al. 2016), may also leverage antibiotic resistance to scavenge biomolecules following post-fire microbial mortality (Willms et al. 2020). It has also been identified as highly connected in desert soil networks (Li et al. 2021), making it well suited to a hub role in the disturbed post-fire environment. It was the only taxon appearing as a hub across three chaparral burned timepoints, suggesting a sustained structural role throughout succession. At 12 months, *Noviherbaspirillum* (Pseudomonadota, module hub) emerged as a keystone, positively associated with the low-abundance saprotrophs *Arsenicitalea* (nitrate reduction; Mu et al. 2016), *Flavitalea* (glycoside hydrolase activity; Kim et al. 2016), *Longimicrobiaceae* (phosphatase activity; Pascual et al. 2016), and *Microvirga* (heat and radiation tolerance; Li et al. 2020). This pattern of a dominant pyrophilous bacterium structuring associations with low-abundance functional degraders may reflect cross-feeding on PyOM, as genomic work has shown that pyrophilous bacteria carry PyOM-degradation genes and that a plasmid from *Noviherbaspirillum soli* has been horizontally transferred into the pyrophilous fungus *Coniochaeta hoffmannii* (Sari et al. 2026), raising the possibility that bacterial-fungal cross-feeding on shared post-fire substrates underlies some of the positive associations we observed. By 40 months, *Candidatus Udaeobacter* remained a hub, now associating with the fungus *Coniochaeta*, while *Gelasinospora* (Ascomycota, connector) linked *Rubrobacter*, *Cladophialophora*, and *Bryobacter* across Module 21. Notably, *Massilia* shifted from a connector hub at 6 months to a peripheral associate of *Candidatus Udaeobacter* by 40 months, suggesting a successional transition in the structural importance of pyrophilous taxa over time. *Mycobacterium* (Actinomycetota) is a connector hub in unburned chaparral networks yet appeared in burned modules as a positive associate of *Candidatus Udaeobacter* at 6 months and a negative associate by 40 months, potentially reflecting competitive dynamics as the post-fire community matures.

### Desert keystones

In contrast to the chaparral, the pyrophilous taxa that increased in abundance following our desert wildfire, including the bacteria *Noviherbaspirillum*, *Tumebacillus*, and *Solirubrobacter* and the fungi *Aspergillus*, *Coniochaeta*, *Penicillium*, and *Naganishia* (Joukhajian et al. 2026) did not emerge as keystones in burned desert networks. However, we identified a putative pyrophilous keystone fungus *Pseudotricharina* (Ascomycota, module hub), as it has been found fruiting post-fire (Cardimona 2023), and is a member of the family Pyronemataceae, which shows phylogenetically conserved fire affinity (Enright et al. 2022).

*Pseudotricharina* appeared as a keystone at 8 months, connected to the bacteria *Pseudonocardia* and fungi *Neophaeococcomyces* and *Inopinatum* in Module 8. The remaining burned desert keystones were generalist bacteria not previously characterized as pyrophilous. The bacterium *wb1-P19* (Pseudomonadota, module hub) was centrally connected at 0.5 months to *Modestobacter* and PMMR1 (*Caulobacter*), and alongside fungal associates including *Acrophialophora* and a Sordariomycetes sp., suggesting cross-kingdom associations from the earliest post-fire timepoint.

Interestingly, *wb1-P19* has been identified as a keystone in cave systems with basic soils in China (Li et al. 2021) and in the nearby Lehman Caves in Nevada (Jones et al. 2024). It possesses methanotroph genes for degrading methanol and formaldehyde, a fire byproduct, as well as a membrane-bound dissimilatory nitrate reductase (NarGHJI), making it functionally well-suited to post-fire substrate utilization (Liao et al. 2021). At 8 months, *CCD24* (Pseudomonadota, module hub) was associated with *Steroidobacter*, *Polycyclovorans*, and *Crossiella*, which later emerged as a module hub. *Rubrobacter* (Actinomycetota, connector) at 12 months connected *Gaiella*, *Blastococcus*, the archaeon *Can. Nitrososphaera*, and the fungus *Aspergillus*. This module may reflect nitrogen-cycling interactions in nutrient-enriched post-fire soils, consistent with elevated ammonium and nitrate detected in burned desert plots (Joukhajian et al. 2026), as both *Rubrobacter* and *Can. Nitrososphaera* were found to carry N cycling genes and increased with N deposition in steppe habitat (Ye et al. 2024). Notably, *Rubrobacter* has been shown to decrease with fire intensity in nearby chaparral and sage scrub environments (Patrick et al. 2024), suggesting its emergence as a desert keystone reflects a context-dependent fire response favored under lower-intensity disturbance. At 36 months, *Crossiella* (Actinomycetota, module hub) associated with the fungi *Conocybe* and *Phaeococcomyces*, as well as *Acidibacter* and *Venturia*, forming cross-kingdom associations at the latest timepoint. *Crossiella* and *wb1-P19* have both been implicated in yellow biofilm formation in cave systems globally (Martin-Pozas et al.

2024), and *Crossiella* may form syntrophic associations with nitrifying bacteria (Martin-Pozas et al. 2022), consistent with its hub role in nutrient-enriched post-fire desert soils. No hub or connector taxa were detected in unburned desert networks, consistent with prokaryotic networks becoming less complex with increasing aridity (Dong et al. 2025).

## Conclusion

Cross-kingdom network analysis revealed that bacterial-fungal interactions were more altered by wildfire than within-kingdom interactions in both systems, with cross-kingdom networks consistently containing more edges than bacterial or fungal networks alone. Across both ecosystems, positive associations increased relative to negative ones in burned plots, meaning cohesion declined, which indicates that wildfire shifted communities toward greater co-occurrence and interdependence regardless of ecosystem type. This shift was detectable across all four post-fire timepoints, demonstrating that the reorganization of bacterial-fungal interactions is a persistent feature of post-fire succession rather than a transient early response. In the desert, where richness was unaffected by fire, network analysis still detected this structural shift, exposing community change missed by traditional diversity metrics. Keystone taxa were system-specific, with pyrophilous bacteria dominating chaparral burned network hubs and generalists dominating the desert, yet bacterial-fungal associations within keystone modules were present at every timepoint in both systems.

As drylands expand and wildfire frequency increases under climate change, understanding how fire reshapes microbial networks and community stability is essential for predicting ecosystem recovery. The persistence of reduced cohesion and altered keystone structure across three years suggests that post-fire dryland soils may remain in a structurally vulnerable state well beyond the window typically assessed in restoration planning. Pyrophilous keystone taxa, particularly those driving cross-kingdom associations with decomposers, represent candidate targets for microbiome-informed restoration strategies, including soil inoculation approaches aimed at restoring competitive network structure and nutrient cycling capacity in severely burned landscapes.

## Acknowledgements

We thank the Cleveland National Forest and the Trabuco Ranger District, including District Ranger Darrel Vance and Emily Fudge, Jeffrey Heys, Lauren Quon, Jacob Rodriguez, and Victoria Stempniewicz for their permitting assistance for the Holy Fire. We thank the Mojave National Preserve including Debra Hughson, Drew Kaiser, and Annasofia Andeski for permitting assistance for the Dome Fire and Tasha La Doux and Jim Andre for assistance at Sweeney Granite Mountain Research Center. We thank Judy Chung, Kobe Luu, Dylan Enright, Aral Greene, Sunny Samear Saroa, Elizah Stephens, Marcos V. Caiafa, and Esbeiry Cordova-Ortiz, for assistance with sample collection and field work. We thank Judy Chung, James Randolph, and Maria Ordoñez for assistance with molecular work and David Lyons at the UCR Environmental Sciences Research Laboratory and Elizah Stephens for help with soil chemical analysis. We thank Joel Sachs and Quinn McFrederick and anonymous reviewers for feedback on the manuscript.

## References

Aanderud ZT, Bahr J, Robinson DM, et al (2019) The Burning of Biocrusts Facilitates the Emergence of a Bare Soil Community of Poorly-Connected Chemoheterotrophic Bacteria With Depressed Ecosystem Services. Front Ecol Evol 7:. 10.3389/fevo.2019.00467

Amarnath K, Narla AV, Pontrelli S, et al (2023) Stress-induced metabolic exchanges between complementary bacterial types underly a dynamic mechanism of inter-species stress resistance. Nat Commun 14:3165. 10.1038/s41467-023-38913-8

Amit G, Bashan A (2023) Top-down identification of keystone taxa in the microbiome. Nat Commun 14:3951. 10.1038/s41467-023-39459-5

Banerjee S, Schlaeppi K, van der Heijden MGA (2018) Keystone taxa as drivers of microbiome structure and functioning. Nat Rev Microbiol 16:567–576. 10.1038/s41579-018-0024-1

Bardgett RD, Freeman C, Ostle NJ (2008) Microbial contributions to climate change through carbon cycle feedbacks. ISME J 2:805–814. 10.1038/ismej.2008.58

Belmok A, Rodrigues-Oliveira T, Lopes FAC, et al (2019) Long-Term Effects of Periodical Fires on Archaeal Communities from Brazilian Cerrado Soils. Archaea 2019:6957210. 10.1155/2019/6957210

Birch JD, Lutz JA, Dickinson MB, et al (2025) Small-scale fire refugia increase soil bacterial and fungal richness and increase community cohesion nine years after fire. Science of The Total Environment 966:178677. 10.1016/j.scitotenv.2025.178677

Blazewicz SJ, Schwartz E, Firestone MK (2014) Growth and death of bacteria and fungi underlie rainfall-induced carbon dioxide pulses from seasonally dried soil. Ecology 95:1162– 1172. 10.1890/13-1031.1

Bolyen E, Rideout JR, Dillon MR, et al (2019) Reproducible, interactive, scalable and extensible microbiome data science using QIIME 2. Nat Biotechnol 37:852–857. 10.1038/s41587-019-0209-9

Brewer TE, Handley KM, Carini P, et al (2016) Genome reduction in an abundant and ubiquitous soil bacterium ‘Candidatus Udaeobacter copiosus.’ Nat Microbiol 2:16198. 10.1038/nmicrobiol.2016.198

Caiafa MV, Nelson AR, Borch T, et al (2023) Distinct fungal and bacterial responses to fire severity and soil depth across a ten-year wildfire chronosequence in beetle-killed lodgepole pine forests. Forest Ecology and Management 544:121160. 10.1016/j.foreco.2023.121160

Caporaso JG, Lauber CL, Walters WA, et al (2011) Global patterns of 16S rRNA diversity at a depth of millions of sequences per sample. PNAS 108:4516–4522. 10.1073/pnas.1000080107

Cardimona W (2023) Pseudotricharina intermedia. In: iNaturalist. https://www.inaturalist.org/observations/153596446. Accessed 11 Nov 2025

Carter MR, Gregorich EG (eds) (2007) Soil Sampling and Methods of Analysis, 2nd edn. CRC Press, Boca Raton

Certini G (2005) Effects of fire on properties of forest soils: a review. Oecologia 143:1–10. 10.1007/s00442-004-1788-8

Certini G, Moya D, Lucas-Borja ME, Mastrolonardo G (2021) The impact of fire on soil-dwelling biota: A review. Forest Ecology and Management 488:118989. 10.1016/j.foreco.2021.118989

Coyte KZ, Schluter J, Foster KR (2015) The ecology of the microbiome: Networks, competition, and stability. Science 350:663–666. 10.1126/science.aad2602

Crowther TW, Hoogen J van den, Wan J, et al (2019) The global soil community and its influence on biogeochemistry. Science 365:. 10.1126/science.aav0550

Csardi MG (2013) Package ‘igraph.’ Last accessed 3:2013

de Vries FT, Griffiths RI, Bailey M, et al (2018) Soil bacterial networks are less stable under drought than fungal networks. Nat Commun 9:3033. 10.1038/s41467-018-05516-7

Deng Y, Jiang Y-H, Yang Y, et al (2012) Molecular ecological network analyses. BMC Bioinformatics 13:113. 10.1186/1471-2105-13-113

Dong L, Bai X, Yu X, et al (2025) Aridity simplified the bacterial network and thus reduced soil multifunctionality in arid and semiarid regions. Plant Soil. 10.1007/s11104-025-07573-6

Dooley SR, Treseder KK (2012) The effect of fire on microbial biomass: a meta-analysis of field studies. Biogeochemistry 109:49–61. 10.1007/s10533-011-9633-8

Enright DJ, Frangioso KM, Isobe K, et al (2022) Mega-fire in redwood tanoak forest reduces bacterial and fungal richness and selects for pyrophilous taxa that are phylogenetically conserved. Molecular Ecology 31:2475–2493. 10.1111/mec.16399

Enright DJ, Veerabahu A, Quaal RJ, et al (2026) Evaluating Best Practices for Isolating Pyrophilous Bacteria and Fungi from Burned Soil. 2024.09.16.612975

Faust K, Lima-Mendez G, Lerat J-S, et al (2015) Cross-biome comparison of microbial association networks. Front Microbiol 6:. 10.3389/fmicb.2015.01200

Feng S, Fu Q (2013) Expansion of global drylands under a warming climate. Atmospheric Chemistry and Physics 13:10081–10094

Fernández-González AJ, Lasa AV, Cobo-Díaz JF, et al (2023) Long-Term Persistence of Three Microbial Wildfire Biomarkers in Forest Soils. Forests 14:1383. 10.3390/f14071383

Ferreira AC, Nobre MF, Moore E, et al (1999) Characterization and radiation resistance of new isolates of Rubrobacter radiotolerans and Rubrobacter xylanophilus. Extremophiles 3:235–238. 10.1007/s007920050121

Fox S, Sikes BA, Brown SP, et al (2022) Fire as a driver of fungal diversity — A synthesis of current knowledge. Mycologia 114:215–241. 10.1080/00275514.2021.2024422

Glassman SI, Randolph JWJ, Saroa SS, et al (2023) Prescribed versus wildfire impacts on exotic plants and soil microbes in California grasslands. Applied Soil Ecology 185:104795. 10.1016/j.apsoil.2022.104795

Guimerà R, Nunes Amaral LA (2005) Functional cartography of complex metabolic networks. Nature 433:895–900. 10.1038/nature03288

Guseva K, Darcy S, Simon E, et al (2022) From diversity to complexity: Microbial networks in soils. Soil Biology and Biochemistry 169:108604. 10.1016/j.soilbio.2022.108604

Hernandez DJ, David AS, Menges ES, et al (2021) Environmental stress destabilizes microbial networks. The ISME Journal 15:1722–1734. 10.1038/s41396-020-00882-x

Herren CM, McMahon KD (2017) Cohesion: a method for quantifying the connectivity of microbial communities. The ISME Journal 11:2426–2438. 10.1038/ismej.2017.91

Hirano H, Takemoto K (2019) Difficulty in inferring microbial community structure based on co-occurrence network approaches. BMC Bioinformatics 20:329. 10.1186/s12859-019-2915-1

Huang J, Yu H, Guan X, et al (2016) Accelerated dryland expansion under climate change. Nature Clim Change 6:166–171. 10.1038/nclimate2837

Jacoby R, Peukert M, Succurro A, et al (2017) The Role of Soil Microorganisms in Plant Mineral Nutrition—Current Knowledge and Future Directions. Front Plant Sci 8:. 10.3389/fpls.2017.01617

Jameson BD, Murdock SA, Ji Q, et al (2023) Network analysis of 16S rRNA sequences suggests microbial keystone taxa contribute to marine N2O cycling. Commun Biol 6:212. 10.1038/s42003-023-04597-5

Jiang M-Z, Zhu H-Z, Zhou N, et al (2022) Droplet microfluidics-based high-throughput bacterial cultivation for validation of taxon pairs in microbial co-occurrence networks. Sci Rep 12:18145. 10.1038/s41598-022-23000-7

Jones DS, Green KM, Brown A, et al (2024) Metagenomic insights into an enigmatic gammaproteobacterium that is important for carbon cycling in cave ecosystems worldwide. 2024.08.23.608578

Jordán F (2009) Keystone species and food webs. Philos Trans R Soc Lond B Biol Sci 364:1733–1741. 10.1098/rstb.2008.0335

Joukhajian A, Barriga MFP, Davis MJ, et al (2026) Mojave Desert microbial communities show high resistance and resilience over three years despite widespread plant mortality following the Dome Fire. fire ecol 22:13. 10.1186/s42408-025-00435-7

Kaiser D, Hughson D (2020) The Dome Fire. Sweeney Granite Mountains Desert Research Center Science Newsletter 20

Kajihara KT, Hynson NA (2024) Networks as tools for defining emergent properties of microbiomes and their stability. Microbiome 12:184. 10.1186/s40168-024-01868-z

Keeley JE (2009) Fire intensity, fire severity and burn severity: a brief review and suggested usage. Int J Wildland Fire 18:116–126. 10.1071/WF07049

Kim S-J, Ahn J-H, Weon H-Y, et al (2016) Flavitalea soli sp. nov. isolated from soil. International Journal of Systematic and Evolutionary Microbiology 66:562–566. 10.1099/ijsem.0.000754

Kõljalg U, Larsson K-H, Abarenkov K, et al (2005) UNITE: a database providing web-based methods for the molecular identification of ectomycorrhizal fungi. New Phytologist 166:1063–1068. 10.1111/j.1469-8137.2005.01376.x

Kurtz ZD, Müller CL, Miraldi ER, et al (2015) Sparse and Compositionally Robust Inference of Microbial Ecological Networks. PLOS Computational Biology 11:e1004226. 10.1371/journal.pcbi.1004226

Lee KK, Kim H, Lee Y-H (2022) Cross-kingdom co-occurrence networks in the plant microbiome: Importance and ecological interpretations. Front Microbiol 13:. 10.3389/fmicb.2022.953300

Li J, Gao R, Chen Y, et al (2020) Isolation and Identification of Microvirga thermotolerans HR1, a Novel Thermo-Tolerant Bacterium, and Comparative Genomics among Microvirga Species. Microorganisms 8:101. 10.3390/microorganisms8010101

Li Q, Song A, Yang H, Müller WEG (2021) Impact of Rocky Desertification Control on Soil Bacterial Community in Karst Graben Basin, Southwestern China. Front Microbiol 12:. 10.3389/fmicb.2021.636405

Li X, Kimball S, Ta P, et al (2024) Understanding post-fire vegetation recovery in southern California ecosystems with the aid of pre-fire observations from long-term monitoring. Journal of Vegetation Science 35:e13308. 10.1111/jvs.13308

Liao J, Wolfe GM, Hannun RA, et al (2021) Formaldehyde evolution in US wildfire plumes during the Fire Influence on Regional to Global Environments and Air Quality experiment (FIREX-AQ). Atmospheric Chemistry and Physics 21:18319–18331. 10.5194/acp-21-18319-2021

Luo W, Wang P, Liu J, Tao J (2025) Microbial keystone taxa and network complexity, rather than diversity, sustain soil multifunctionality along an elevational gradient in a subtropical karst mountain. CATENA 256:109115. 10.1016/j.catena.2025.109115

Martin-Pozas T, Cuezva S, Fernandez-Cortes A, et al (2022) Role of subterranean microbiota in the carbon cycle and greenhouse gas dynamics. Science of The Total Environment 831:154921. 10.1016/j.scitotenv.2022.154921

Martin-Pozas T, Jurado V, Fernandez-Cortes A, et al (2024) Bacterial communities forming yellow biofilms in different cave types share a common core. Science of The Total Environment 956:177263. 10.1016/j.scitotenv.2024.177263

Matchado MS, Lauber M, Reitmeier S, et al (2021) Network analysis methods for studying microbial communities: A mini review. Computational and Structural Biotechnology Journal 19:2687–2698. 10.1016/j.csbj.2021.05.001

Meinshausen N, Bühlmann P (2006) High-dimensional graphs and variable selection with the Lasso. Annals of Statistics 34:

Morriën E, Hannula SE, Snoek LB, et al (2017) Soil networks become more connected and take up more carbon as nature restoration progresses. Nat Commun 8:14349. 10.1038/ncomms14349

Mu Y, Zhou L, Zeng X-C, et al (2016) Arsenicitalea aurantiaca gen. nov., sp. nov., a new member of the family Hyphomicrobiaceae, isolated from high-arsenic sediment. International Journal of Systematic and Evolutionary Microbiology 66:5478–5484. 10.1099/ijsem.0.001543

Nelson AR, Narrowe AB, Rhoades CC, et al (2022) Wildfire-dependent changes in soil microbiome diversity and function. Nat Microbiol 7:1419–1430. 10.1038/s41564-022-01203-y

Nelson AR, Rhoades CC, Fegel TS, et al (2024) Wildfire impact on soil microbiome life history traits and roles in ecosystem carbon cycling. ISME Commun 4:ycae108. 10.1093/ismeco/ycae108

Pascual J, García-López M, Bills GF, Genilloud O (2016) Longimicrobium terrae gen. nov., sp. nov., an oligotrophic bacterium of the under-represented phylum Gemmatimonadetes isolated through a system of miniaturized diffusion chambers. International Journal of Systematic and Evolutionary Microbiology 66:1976–1985. 10.1099/ijsem.0.000974

Patrick TA, Maguire LW, Espin J, Gardner CM (2024) Wildfire Impacts on Soil Microbiomes: Potential for Disruptions to Nitrogen-Cycling Bacteria. Environmental Engineering Science 41:337–346. 10.1089/ees.2024.0074

Pausas JG, Keeley JE (2021) Wildfires and global change. Frontiers in Ecology and the Environment 19:387–395. 10.1002/fee.2359

Pérez-Valera E, Goberna M, Faust K, et al (2017) Fire modifies the phylogenetic structure of soil bacterial co-occurrence networks. Environmental Microbiology 19:317–327. 10.1111/1462-2920.13609

Pérez-Valera E, Verdú M, Navarro-Cano JA, Goberna M (2018) Resilience to fire of phylogenetic diversity across biological domains. Molecular Ecology 27:2896–2908. 10.1111/mec.14729

Power ME, Tilman D, Estes JA, et al (1996) Challenges in the quest for keystones: identifying keystone species is difficult—but essential to understanding how loss of species will affect ecosystems. BioScience 46:609–620

Pressler Y, Foster EJ, Moore JC, Cotrufo MF (2017) Coupled biochar amendment and limited irrigation strategies do not affect a degraded soil food web in a maize agroecosystem, compared to the native grassland. GCB Bioenergy 9:1344–1355. 10.1111/gcbb.12429

Pressler Y, Moore JC, Cotrufo MF (2019) Belowground community responses to fire: meta-analysis reveals contrasting responses of soil microorganisms and mesofauna. Oikos 128:309–327. 10.1111/oik.05738

Pulido Barriga MF, Kuhn AN, Homyak PM, et al (2025) Chaparral wildfire shifts the functional potential for soil pyrogenic organic matter and nitrogen cycling. Soil Biology and Biochemistry 211:109985. 10.1016/j.soilbio.2025.109985

Pulido-Chavez MF (2023) Measuring the Succession, Functions and Resilience of Soil Microbes After a Chaparral Wildfire. Ph.D., University of California, Riverside

Pulido-Chavez MF, Randolph JWJ, Zalman C, et al (2023) Rapid bacterial and fungal successional dynamics in first year after chaparral wildfire. Molecular Ecology 32:1685– 1707. 10.1111/mec.16835

Quast C, Pruesse E, Yilmaz P, et al (2013) The SILVA ribosomal RNA gene database project: improved data processing and web-based tools. Nucleic Acids Research 41:D590–D596. 10.1093/nar/gks1219

R Core Team (2024) R: A language and environment for statistical computing. https://www.R-project.org/. Accessed 22 Aug 2021

Rawstern AH, Hernandez DJ, Afkhami ME (2025) Central Taxa Are Keystone Microbes During Early Succession. Ecology Letters 28:e70031. 10.1111/ele.70031

Ren J, Hanan EJ, Greene A, et al (2024) Simulating the Role of Biogeochemical Hotspots in Driving Nitrogen Export From Dryland Watersheds. Water Resources Research 60:e2023WR036008. 10.1029/2023WR036008

Röttjers L, Faust K (2018) From hairballs to hypotheses–biological insights from microbial networks. FEMS Microbiology Reviews 42:761–780. 10.1093/femsre/fuy030

Sari E, Enright DJ, Ordonez M, et al (2025) Strain-Level Adaptation of Pyrophilous Bacteria through Gene Fragmentation and Horizontal Gene Transfer. 2025.11.11.687939

Sari E, Enright DJ, Ordoñez ME, et al (2026) Gene duplication, horizontal gene transfer, and trait trade-offs drive evolution of postfire resource acquisition in pyrophilous fungi. Proceedings of the National Academy of Sciences 123:e2519152123. 10.1073/pnas.2519152123

Schlesinger WH, Reynolds JF, Cunningham GL, et al (1990) Biological feedbacks in global desertification. Science 247:1043–1048

Soil Survey Staff (2024) Web Soil Survey. https://websoilsurvey.nrcs.usda.gov/. Accessed 22 June 2026

Soil Survey Staff (2015) MINEHART Soil Series. In: Web Soil Survey. https://soilseries.sc.egov.usda.gov/OSD_Docs/M/MINEHART.html. Accessed 9 June 2025

Soria R, Tortosa A, Rodríguez-Berbel N, et al (2023) Short-Term Response of Soil Bacterial Communities after Prescribed Fires in Semi-Arid Mediterranean Forests. Fire 6:145. 10.3390/fire6040145

Stanton RL, Nusink BC, Cass KL, et al (2023) Fire frequency effects on plant community characteristics in the Great Basin and Mojave deserts of North America. Fire Ecology 19:60. 10.1186/s42408-023-00222-2

Stephens EZ, Pulido Barriga MF, Greene AC, et al (2026) Wildfire alters nitrogen cycling to increase soil emissions of nitric oxide (NO) and the heterogeneity of nitrous oxide (N2O) in California chaparral. Geoderma 469:117807. 10.1016/j.geoderma.2026.117807

Su W, Tang C, Lin J, et al (2022) Recovery patterns of soil bacterial and fungal communities in Chinese boreal forests along a fire chronosequence. Science of The Total Environment 805:150372. 10.1016/j.scitotenv.2021.150372

Taylor DL, Walters WA, Lennon NJ, et al (2016) Accurate Estimation of Fungal Diversity and Abundance through Improved Lineage-Specific Primers Optimized for Illumina Amplicon Sequencing. Appl Environ Microbiol 82:7217–7226. 10.1128/AEM.02576-16

Vega-Cofre MV, Williams W, Song Y, et al (2023) Effects of grazing and fire management on rangeland soil and biocrust microbiomes. Ecological Indicators 148:110094. 10.1016/j.ecolind.2023.110094

Velez P, Espinosa-Asuar L, Figueroa M, et al (2018) Nutrient Dependent Cross-Kingdom Interactions: Fungi and Bacteria From an Oligotrophic Desert Oasis. Front Microbiol 9:. 10.3389/fmicb.2018.01755

Wang X, Liang C, Mao J, et al (2023) Microbial keystone taxa drive succession of plant residue chemistry. ISME J 17:748–757. 10.1038/s41396-023-01384-2

Warcup JH (1990) Occurrence of ectomycorrhizal and saprophytic discomycetes after a wild fire in a eucalypt forest. Mycological Research 94:1065–1069. 10.1016/S0953-7562(09)81334-8

Whitman T, Whitman E, Woolet J, et al (2019) Soil bacterial and fungal response to wildfires in the Canadian boreal forest across a burn severity gradient. Soil Biology and Biochemistry 138:107571. 10.1016/j.soilbio.2019.107571

Whitman T, Woolet J, Sikora M, et al (2022) Resilience in soil bacterial communities of the boreal forest from one to five years after wildfire across a severity gradient. Soil Biology and Biochemistry 172:108755. 10.1016/j.soilbio.2022.108755

Whitman T, Woolet J, Sikora MC, et al (2025) Resilience not yet apparent in soil fungal communities of the boreal forest from one to five years after wildfire across a severity gradient. 2025.03.29.646032

Willms IM, Rudolph AY, Göschel I, et al (2020) Globally Abundant “Candidatus Udaeobacter” Benefits from Release of Antibiotics in Soil and Potentially Performs Trace Gas Scavenging. mSphere 5:10.1128/msphere.00186-20. 10.1128/msphere.00186-20

Woolet J, Whitman E, Parisien M-A, et al (2022) Effects of short-interval reburns in the boreal forest on soil bacterial communities compared to long-interval reburns. FEMS Microbiol Ecol 98:fiac069. 10.1093/femsec/fiac069

Wu D, Bai H, Zhao C, et al (2023) The characteristics of soil microbial co-occurrence networks across a high-latitude forested wetland ecotone in China. Front Microbiol 14:. 10.3389/fmicb.2023.1160683

Xu Q, Vandenkoornhuyse P, Li L, et al (2022) Microbial generalists and specialists differently contribute to the community diversity in farmland soils. Journal of Advanced Research 40:17–27. 10.1016/j.jare.2021.12.003

Yang M, Luo X, Cai Y, et al (2024) Effect of fire and post-fire management on soil microbial communities in a lower subtropical forest ecosystem after a mountain fire. Journal of Environmental Management 351:119885. 10.1016/j.jenvman.2023.119885

Yang T, Tedersoo L, Liu X, et al (2022) Fungi stabilize multi-kingdom community in a high elevation timberline ecosystem. iMeta 1:e49. 10.1002/imt2.49

Ye H, Zhao Y, He S, et al (2024) Metagenomics reveals the response of desert steppe microbial communities and carbon-nitrogen cycling functional genes to nitrogen deposition. Front Microbiol 15:. 10.3389/fmicb.2024.1369196

Zeng N, Yoon J (2009) Expansion of the world’s deserts due to vegetation-albedo feedback under global warming. Geophysical Research Letters 36:. 10.1029/2009GL039699

Zhang Y, Du Y, Mu Z, et al (2025) Response of Soil Microbial Communities in Extreme Arid Deserts to Different Long-Term Management Methods. Forests 16:306. 10.3390/f16020306

