## Supplemental Tables and Figures for "Microbial association networks reveal hidden keystone taxa and cross-kingdom interactions after dryland wildfires"

### Supplementary Table of Contents

Table S1. Chaparral network metrics

Table S2. Desert network metrics

Table S3. Keystone taxa in chaparral and desert networks

Figure S1. Soil edaphic characteristics

Figure S2. Bacteria and Fungi OTU richness

Figure S3. Network-wide positive edge percentage

Figure S4. Sub-sampled cohesion in cross-kingdom networks

Figure S5. Chaparral unburned community keystone taxa

### Supplementary Tables

Table S1. Chaparral network metrics per timepoint in 16S, ITS2, and cross-kingdom network analyses. Nodes indicate genera which passed our filters, and edges are positive or negative associations above 1e-8.

| **Timepoint** | **Treatment** | **Type** | **Nodes** | **Total Edges** | **Positive Edges** | **Negative Edges** | **Modularity** | **16S-ITS2 Samples** |
| --- | --- | --- | --- | --- | --- | --- | --- | --- |
| 0.5 Months | Burned | 16S | 227 | 63 | 54 | 9 | 0.872 | 21-0 |
| 6 Months | Burned | 16S | 164 | 40 | 33 | 7 | 0.843 | 24-0 |
| 12 Months | Burned | 16S | 231 | 79 | 67 | 12 | 0.779 | 24-0 |
| 40 Months | Burned | 16S | 262 | 74 | 53 | 21 | 0.897 | 24-0 |
| 0.5 Months | Unburned | 16S | 308 | 79 | 64 | 15 | 0.962 | 12-0 |
| 6 Months | Unburned | 16S | 391 | 122 | 97 | 25 | 0.979 | 12-0 |
| 12 Months | Unburned | 16S | 289 | 49 | 39 | 10 | 0.962 | 12-0 |
| 40 Months | Unburned | 16S | 300 | 66 | 50 | 16 | 0.969 | 12-0 |
| 0.5 Months | Burned | Cross | 300 | 76 | 53 | 23 | 0.937 | 21-21 |
| 6 Months | Burned | Cross | 260 | 94 | 73 | 21 | 0.822 | 24-24 |
| 12 Months | Burned | Cross | 311 | 130 | 104 | 26 | 0.818 | 24-24 |
| 40 Months | Burned | Cross | 358 | 146 | 115 | 31 | 0.898 | 24-24 |
| 0.5 Months | Unburned | Cross | 593 | 429 | 313 | 116 | 0.939 | 12-12 |
| 6 Months | Unburned | Cross | 668 | 576 | 361 | 215 | 0.936 | 12-12 |
| 12 Months | Unburned | Cross | 512 | 273 | 197 | 76 | 0.968 | 12-12 |
| 40 Months | Unburned | Cross | 532 | 261 | 182 | 79 | 0.976 | 12-12 |
| 0.5 Months | Burned | ITS2 | 87 | 3 | 3 | 0 | 0.667 | 0-23 |
| 6 Months | Burned | ITS2 | 96 | 17 | 17 | 0 | 0.746 | 0-24 |
| 12 Months | Burned | ITS2 | 80 | 7 | 6 | 1 | 0.857 | 0-24 |
| 40 Months | Burned | ITS2 | 96 | 13 | 12 | 1 | 0.852 | 0-24 |
| 0.5 Months | Unburned | ITS2 | 285 | 66 | 56 | 10 | 0.951 | 0-12 |
| 6 Months | Unburned | ITS2 | 277 | 60 | 57 | 3 | 0.96 | 0-12 |
| 12 Months | Unburned | ITS2 | 223 | 37 | 35 | 2 | 0.948 | 0-12 |
| 40 Months | Unburned | ITS2 | 232 | 36 | 32 | 4 | 0.929 | 0-12 |

Table S2. Desert network metrics per timepoint in 16S, ITS2, and cross-kingdom network analyses. Nodes indicate genera which passed our filters, and edges are positive or negative associations above 1e-8.

| **Timepoint** | **Treatment** | **Type** | **Nodes** | **Total Edges** | **Positive Edges** | **Negative Edges** | **Modularity** | **16S-ITS2 Samples** |
| --- | --- | --- | --- | --- | --- | --- | --- | --- |
| 0.5 Months | Burned | 16S | 265 | 74 | 52 | 22 | 0.865 | 24-0 |
| 8 Months | Burned | 16S | 249 | 58 | 42 | 16 | 0.92 | 24-0 |
| 12 Months | Burned | 16S | 243 | 59 | 43 | 16 | 0.893 | 24-0 |
| 36 Months | Burned | 16S | 270 | 57 | 47 | 10 | 0.859 | 24-0 |
| 0.5 Months | Unburned | 16S | 271 | 40 | 33 | 7 | 0.95 | 12-0 |
| 8 Months | Unburned | 16S | 227 | 27 | 27 | 0 | 0.936 | 12-0 |
| 12 Months | Unburned | 16S | 221 | 18 | 13 | 5 | 0.944 | 12-0 |
| 36 Months | Unburned | 16S | 230 | 20 | 16 | 4 | 0.935 | 12-0 |
| 0.5 Months | Burned | Cross | 420 | 170 | 123 | 47 | 0.897 | 24-24 |
| 8 Months | Burned | Cross | 381 | 146 | 95 | 51 | 0.95 | 24-24 |
| 12 Months | Burned | Cross | 377 | 139 | 96 | 43 | 0.919 | 24-24 |
| 36 Months | Burned | Cross | 431 | 166 | 118 | 48 | 0.948 | 24-24 |
| 0.5 Months | Unburned | Cross | 429 | 117 | 86 | 31 | 0.975 | 12-12 |
| 8 Months | Unburned | Cross | 349 | 62 | 53 | 9 | 0.961 | 12-12 |
| 12 Months | Unburned | Cross | 364 | 86 | 59 | 27 | 0.966 | 12-12 |
| 36 Months | Unburned | Cross | 387 | 66 | 45 | 21 | 0.974 | 12-12 |
| 0.5 Months | Burned | ITS2 | 155 | 16 | 12 | 4 | 0.867 | 0-24 |
| 8 Months | Burned | ITS2 | 132 | 15 | 14 | 1 | 0.898 | 0-24 |
| 12 Months | Burned | ITS2 | 134 | 15 | 11 | 4 | 0.871 | 0-24 |
| 36 Months | Burned | ITS2 | 161 | 28 | 19 | 9 | 0.921 | 0-24 |
| 0.5 Months | Unburned | ITS2 | 158 | 15 | 15 | 0 | 0.88 | 0-12 |
| 8 Months | Unburned | ITS2 | 122 | 12 | 12 | 0 | 0.819 | 0-12 |
| 12 Months | Unburned | ITS2 | 143 | 17 | 16 | 1 | 0.851 | 0-12 |
| 36 Months | Unburned | ITS2 | 157 | 8 | 6 | 2 | 0.781 | 0-12 |

Table S3. Keystone taxa with either Pi > 0.62 or Zi >= 2.5 in chaparral and desert networks. Note that unburned keystones were only present in the chaparral.

| Keystone | Kingdom | Type | Module | Zi | Pi | Treatment | Timepoint |
| --- | --- | --- | --- | --- | --- | --- | --- |
| *0319-7L14* | Bacteria | Desert | 45 | 0.378 | 0.640 | Burned | 0.5 Months |
| *Rubrobacter* | Bacteria | Desert | 12 | -0.829 | 0.667 | Burned | 0.5 Months |
| *CCD24* | Bacteria | Desert | 9 | 2.536 | 0.000 | Burned | 8 Months |
| *Pseudotricharina* | Fungi | Desert | 8 | 2.959 | 0.000 | Burned | 8 Months |
| *Rubrobacter* | Bacteria | Desert | 31 | -0.132 | 0.625 | Burned | 12 Months |
| *Crossiella* | Bacteria | Desert | 57 | 2.784 | 0.000 | Burned | 36 Months |
| *Roseiarcus* | Bacteria | Chaparral | 6 | -0.178 | 0.625 | Unburned | 0.5 Months |
| *Mycobacterium* | Bacteria | Chaparral | 6 | -0.178 | 0.625 | Unburned | 0.5 Months |
| *Jatrophihabitans* | Bacteria | Chaparral | 20 | 2.964 | 0.000 | Unburned | 0.5 Months |
| *Novosphingobium* | Bacteria | Chaparral | 27 | -0.907 | 0.667 | Unburned | 6 Months |
| *B12-WMSP1* | Bacteria | Chaparral | 23 | -0.238 | 0.625 | Unburned | 6 Months |
| *Lacunisphaera* | Bacteria | Chaparral | 5 | 2.547 | 0.000 | Unburned | 6 Months |
| *Myxococcus* | Bacteria | Chaparral | 12 | 2.595 | 0.000 | Unburned | 6 Months |
| *Afipia* | Bacteria | Chaparral | 3 | -0.869 | 0.667 | Unburned | 6 Months |
| *Bordetella* | Bacteria | Chaparral | 27 | 2.722 | 0.000 | Unburned | 6 Months |
| *Exophiala* | Fungi | Chaparral | 23 | 3.015 | 0.000 | Unburned | 6 Months |
| *Ceriporia* | Fungi | Chaparral | 27 | 2.722 | 0.000 | Unburned | 6 Months |
| *Massilia* | Bacteria | Chaparral | 1 | 0.185 | 0.625 | Burned | 6 Months |
| *Can. Udaeobacter* | Bacteria | Chaparral | 2 | -0.232 | 0.640 | Burned | 6 Months |
| *Inocybe* | Fungi | Chaparral | 1 | -0.833 | 0.667 | Burned | 6 Months |
| *Chaetosphaeronema* | Fungi | Chaparral | 41 | 2.598 | 0.000 | Unburned | 12 Months |
| *Dendrophoma* | Fungi | Chaparral | 33 | 3.430 | 0.000 | Unburned | 12 Months |
| *Noviherbaspirillum* | Bacteria | Chaparral | 10 | 2.287 | 0.656 | Burned | 12 Months |
| *67-14* | Bacteria | Chaparral | 33 | 2.828 | 0.000 | Burned | 12 Months |
| *Can. Udaeobacter* | Bacteria | Chaparral | 11 | 3.121 | 0.514 | Burned | 12 Months |
| *Can. Udaeobacter* | Bacteria | Chaparral | 16 | 4.092 | 0.165 | Burned | 40 Months |
| *Gelasinospora* | Fungi | Chaparral | 21 | -0.942 | 0.667 | Burned | 40 Months |

### Supplemental Figures

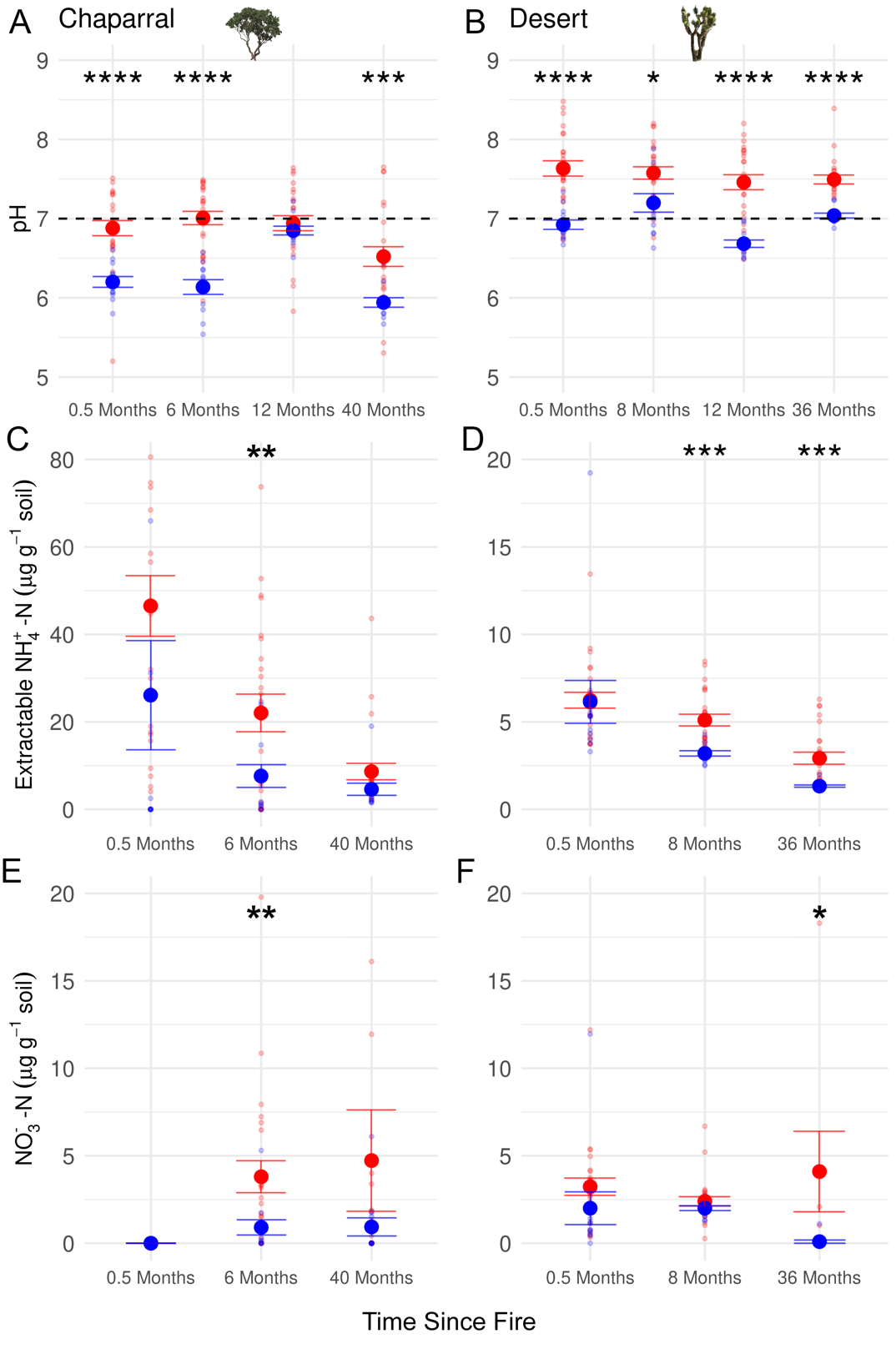

Figure S1. Fire impacts on soil by the Holy Fire (chaparral) and the Dome Fire (desert). Data for the chaparral site was obtained from Stephens et al (2026) and desert from (Joukhajian et al. 2026). Red and blue points indicate means of burned and unburned plots with standard error bars. We measured pH at every timepoint, and measured ammonium and nitrate through KCl extractions at select timepoints. Asterisks indicate significant t-tests (ns = p ≥ 0.05, * = p < 0.05, ** = p < 0.01, *** = p < 0.001, and **** = p < 0.0001). Dots indicate the per treatment mean and bars indicate standard error (burned n=24, unburned n=12).

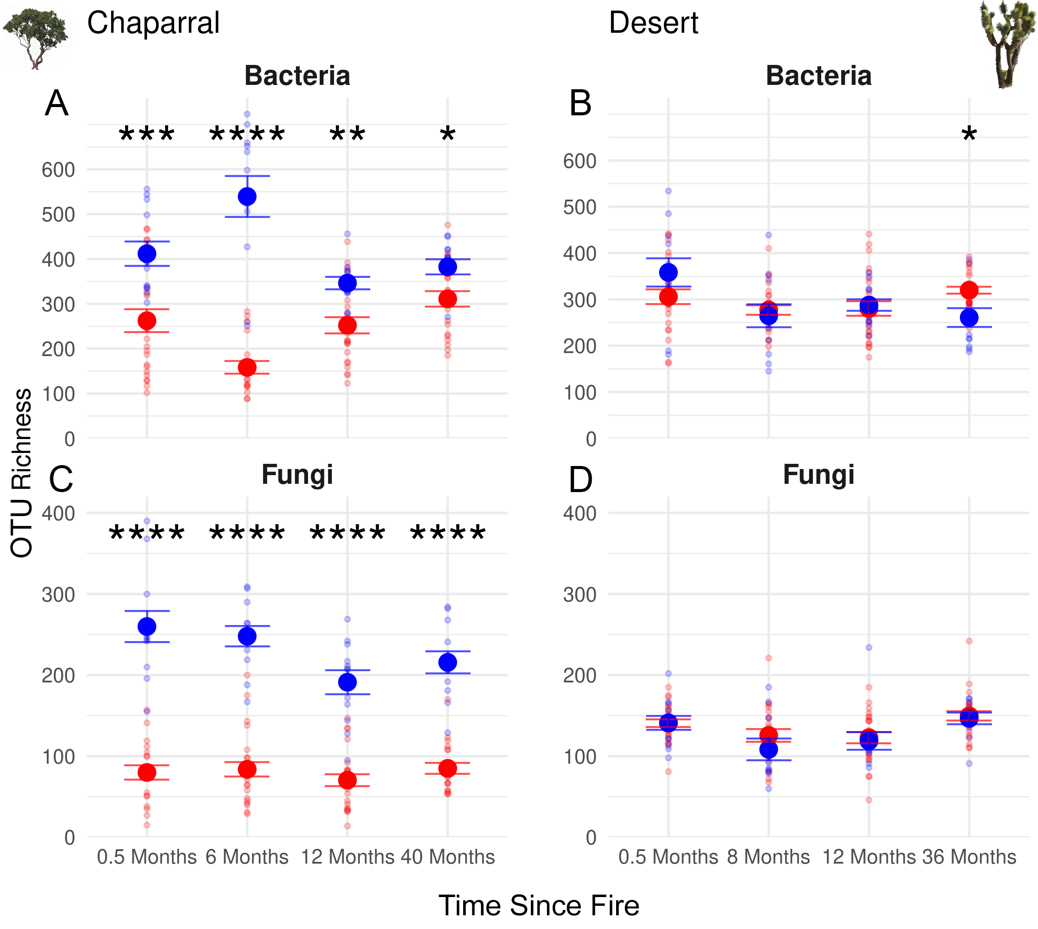

Figure S2. Richness of Operational Taxonomic Units (OTUs) in bacteria as determined by 16S sequencing from A) the chaparral and B) desert. OTU richness in fungi as determined by ITS2 sequencing from C) the chaparral and D) desert. Asterisks indicate significant t tests (ns = p ≥ 0.05, * = p < 0.05, ** = p < 0.01, *** = p < 0.001, and **** = p < 0.0001). Dots indicate the per treatment mean richness and bars indicate standard error (burned n=24, unburned n=12). Data for the chaparral site was obtained from Pulido-Chavez et al. (2023) and desert from Joukhajian et al. (2026).

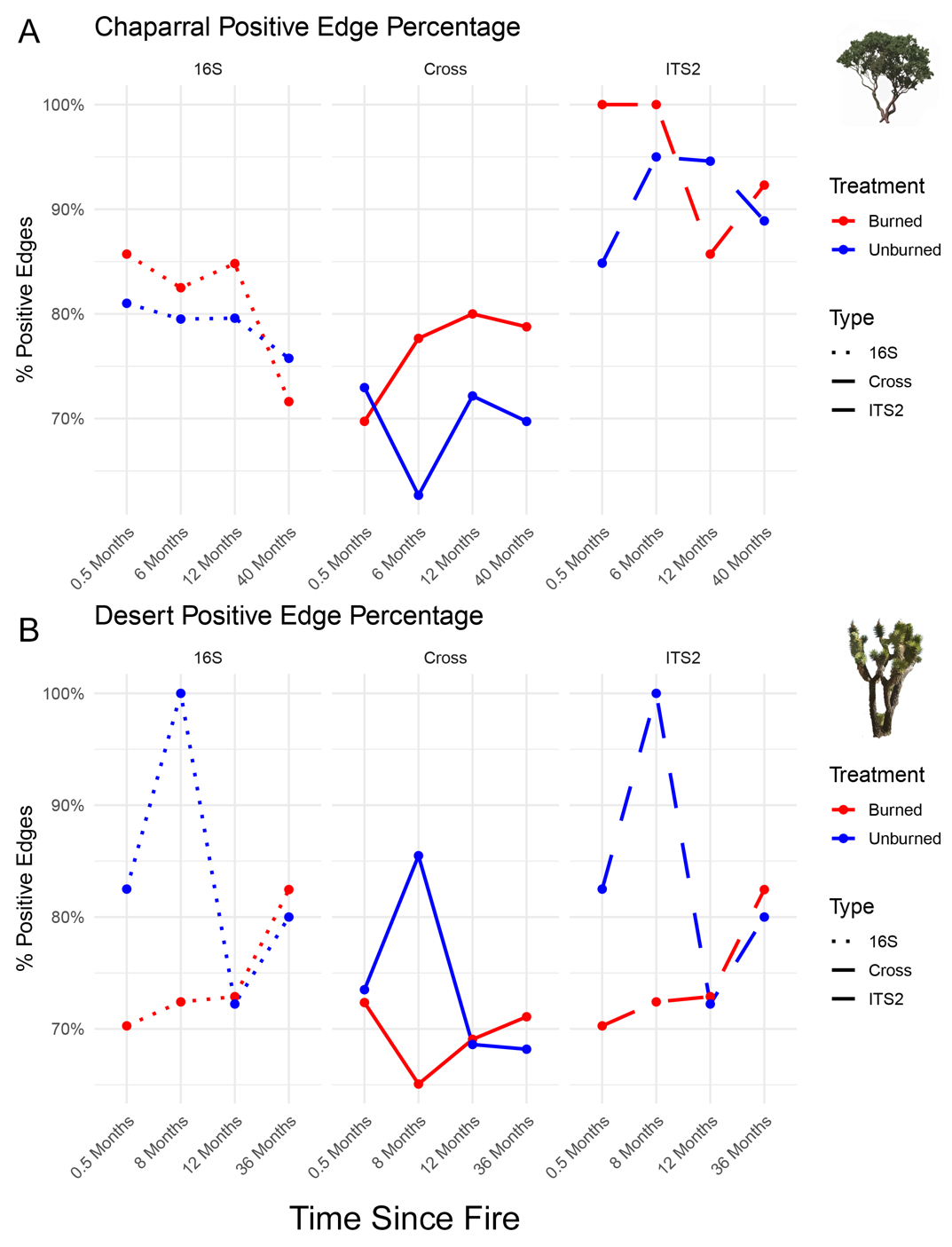

Figure S3. Positive edge percentage of single and cross-kingdom networks from A) chaparral and B) desert burned and unburned plots.

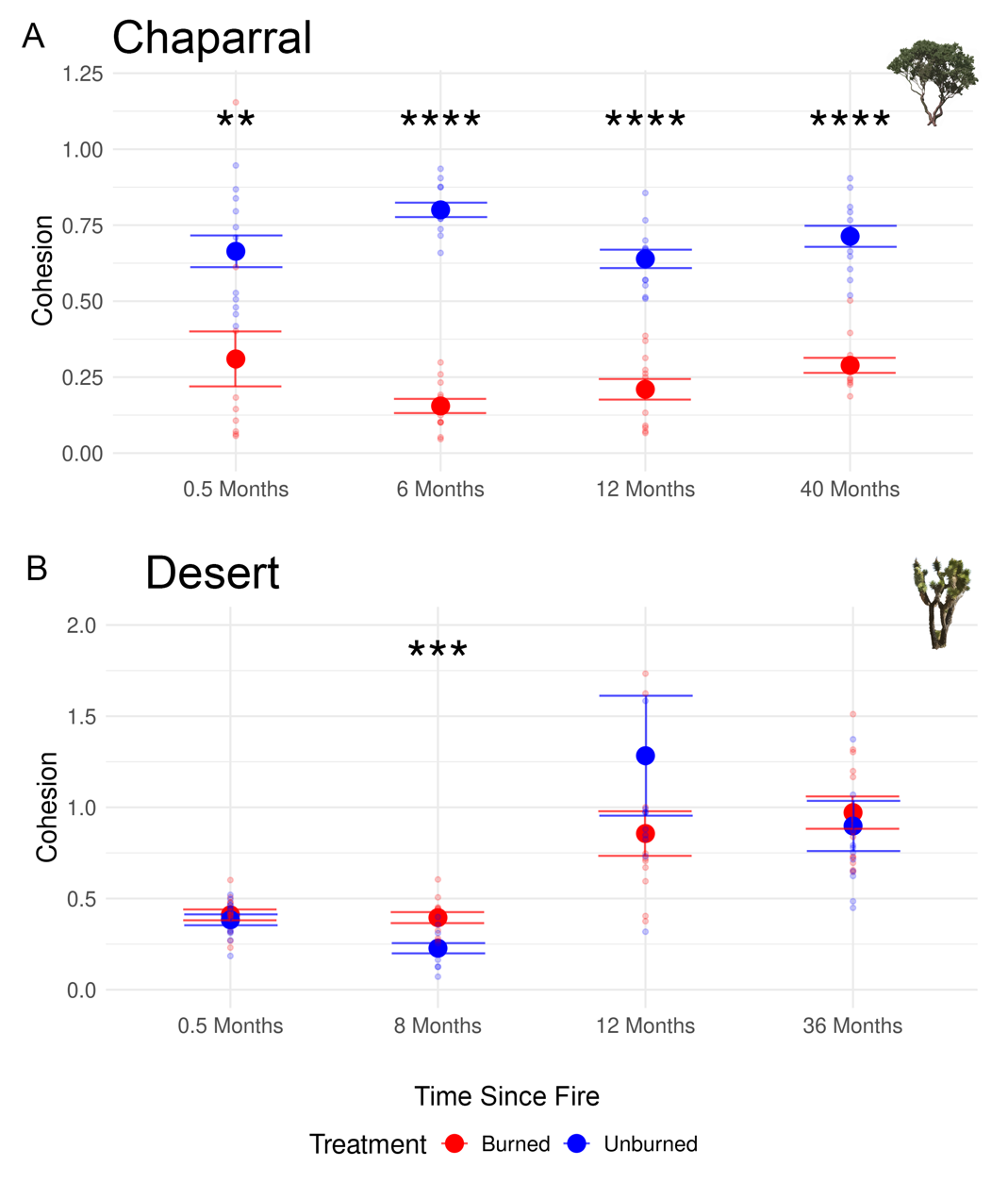

Figure S4. Cross-kingdom cohesion of (A) chaparral and (B) desert networks after subsampling burned networks to match the number of unburned samples. Burned networks were constructed with n = 12 samples, randomly selected from a pool of 24 burned samples per timepoint. Asterisks indicate significant t-tests (ns = p ≥ 0.05, * = p < 0.05, ** = p < 0.01, *** = p < 0.001, and **** = p < 0.0001).

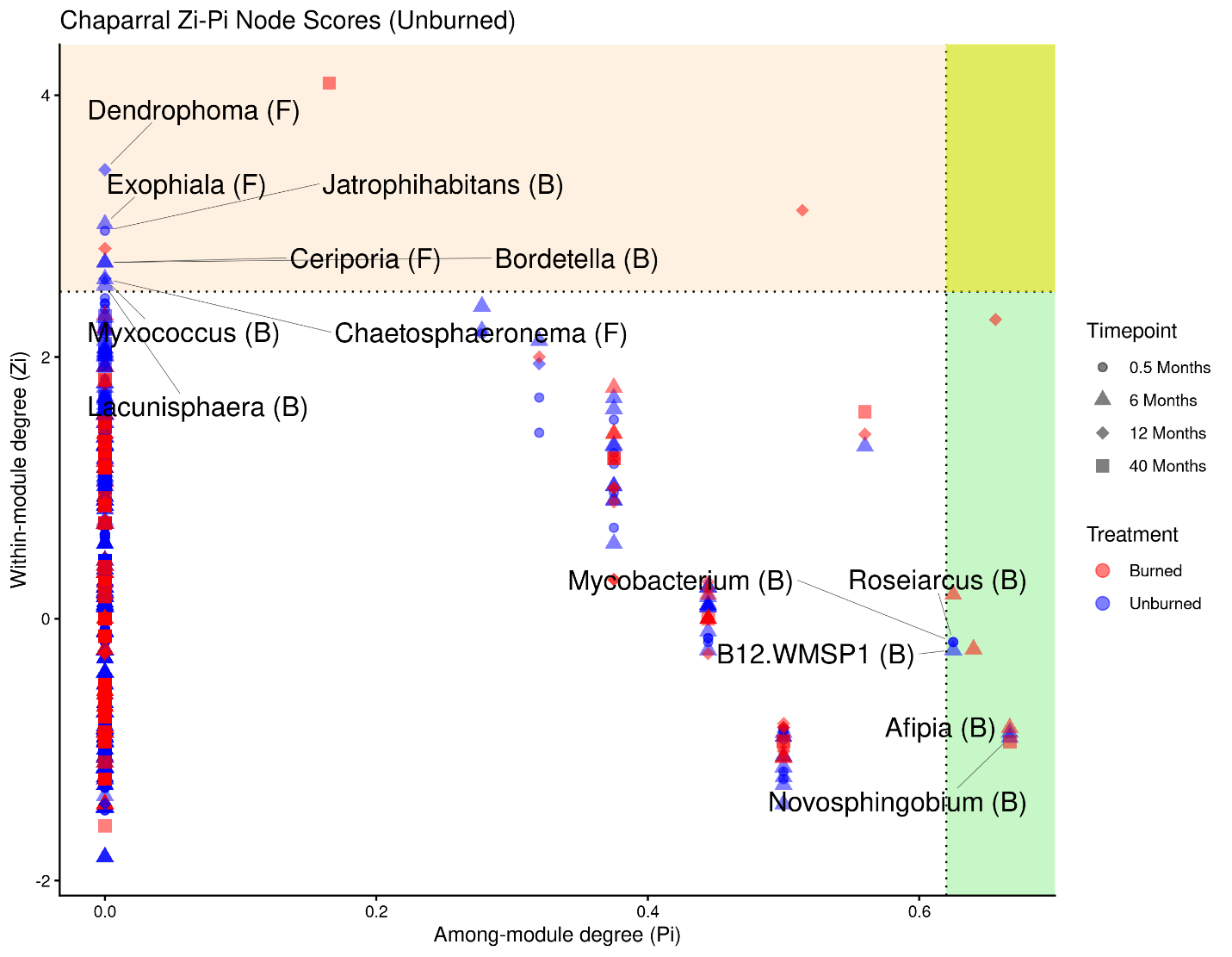

Figure S5. Zi-Pi plots of cross-kingdom networks from unburned chaparral networks at all timepoints. Shapes indicate timepoints, and colors indicate treatment (red for burned and blue for unburned). Peripheral taxa in the white region are below the threshold of being module hubs or connector nodes. Peripheral taxa have few connections (Zi < 2.5 and Pi < 0.62), module hubs have in-module connectivity or many edges (Zi > 2.5, Pi < 0.62; **tan region**), connector nodes have many edges across modules (Zi < 2.5, Pi > 0.62; **green region**), and network hubs are highly connected both within and across modules (Zi >2.5, Pi > 0.62; intersection; **yellow region**).
